# TAOK1 regulates chemo- and radiosensitivity in BRCA1/2-deficient tumors

**DOI:** 10.1101/2025.10.14.682031

**Authors:** Lea Lingg, Merve Mutlu, Arunasalam Naguleswaran, Ismar Klebic, Ewa Gogola, Dilara Akhoundova Sanoyan, Pascale Martine Schürch, Raviprasad Kuthethur, Carmen Fonseca, Simone de Brot, Marieke Everts, Martín González-Fernández, Saskia Hussung, Erik Vassella, Marcel A. T. M. van Vugt, Arnab Ray Chaudhuri, Jos Jonkers, Sven Rottenberg, Paola Francica

## Abstract

BRCA1/2-deficient cells are hypersensitive to replication stress- and DNA-damage-inducing agents such as poly(ADP-ribose) polymerase inhibitors (PARPi), platinum drugs, and ionizing radiation (IR), largely due to impaired homologous recombination repair and replication fork (RF) protection. The precise mechanisms underlying RF vulnerability in the absence of BRCA1/2 remain incompletely understood, however. Here, we identify Thousand And One Amino Acid Kinase 1 (TAOK1) as a novel regulator of RF dynamics that sensitizes BRCA1/2-deficient cells to both PARPi and IR. Unlike established DNA repair factors, such as PARG or the 53BP1-RIF1-shieldin-CST pathway, which mediate PARPi sensitivity while protecting cells against IR, TAOK1 promotes RF degradation and suppresses post-replicative damage repair. TAOK1 depletion stabilizes RFs and reduces replication-associated DNA damage in BRCA1/2-deficient cells, thereby conferring therapy resistance. Interestingly, we found that TAOK1 functions in RF regulation to be independent of its kinase activity. Instead, it interacts with PCNA and regulates ISG15 levels and thereby affects RF stability. Our findings reveal TAOK1 as a non-canonical mediator of therapy sensitivity in BRCA1/2-deficient tumors.

**Teaser:** TAOK1 controls replication fork stability through PCNA and ISG15, affecting therapy response in BRCA1/2-deficient cancers.

## Introduction

Homologous-recombination-deficient (HRD) cancers, in particular those lacking functional BRCA1 or BRCA2, are therapeutically vulnerable to DNA damage-inducing agents. In BRCA1/2-deficient cells, the inability to repair DNA double strand breaks (DSBs) by homologous recombination (HR) leads to genomic instability and increased replication stress (*1*). Therefore, treatments inducing extensive DNA damage, such as radiotherapy (RT), poly(ADP) ribose polymerase inhibitors (PARPi) or platinum agents, are widely used in HRD cancers (*2*, *3*). These agents induce replication stress by generating single strand breaks (SSBs), trapped PARP-DNA complexes, transcription-replication conflicts, or DNA adducts, which eventually stall replication forks (RFs) and trigger cell death (*4–8*). It was previously shown that BRCA1/2-deficient cells are particularly sensitive to DNA damage-induced replication stress due to their susceptibility to degradation of stalled RFs (*9–11*). Although in normal cells the main mechanism of this sensitivity appears to be the inability to repair the subsequent fork collapse by HR (*12*), BRCA1/2-deficient cancers can also acquire resistance to DNA damage-inducing therapy independently of HR restoration (*13*). Restoration of RF stability, by depletion of PTIP, CHD4, PARP1 (*11*), H2AX (*14*), or an impaired DNA prereplication complex (*15*), also results in chemoresistance.

A major regulator of processing damage at replication sites is the DNA sliding clamp PCNA, which requires post-translational modifications (PTMs) to activate down-stream pathways. Mono- and polyubiquitination of PCNA activate DNA damage tolerance (DDT) pathways, involving either translesion synthesis (TLS) or template switching (TS) and allow cells to bypass replication lesions (*16*, *17*). However, TLS polymerases are error-prone, contributing to an increased mutational burden in BRCA1/2-deficient cells, which is characterized by an accumulation of single-nucleotide variations (SNV) (*18*). Ubiquitinated PCNA was further shown to be implicated in RF stability in BRCA1/2-deficient cells (*19*). An opposing role to ubiquitin-mediated PCNA regulation involves ISG15-dependent modification of PCNA. Specifically, ISGylation of PCNA leads to the removal of ubiquitin and inhibits TLS (*20*). Additionally, ISG15 was shown to be essential for RF restart in cells that undergo replication stress (*21*). Not only PTMs on PCNA but also modification of other factors recruited to RFs thus seem crucial for processing replication-related DNA damage.

A tight regulation of RF integrity and replication-associated DNA repair is clearly emerging as one of the crucial factors influencing the response to DNA damage-inducing agents in BRCA1/2-deficient models (*13*). However, the underlying molecular mechanisms are still ill-defined and there is an urgent need to shed light on the regulation of RF integrity in relation to DNA damage tolerance in HR-deficient cells. The response of HRD tumors to DNA damage-inducing therapies, such as PARPi and ionizing radiation (IR), is influenced by distinct and sometimes opposing molecular mechanisms. Proteins including 53BP1 (*22*), RIF1 (*23*), REV7 (*24*), the shieldin complex (*25*) and CST complexes (*26*) are known to mediate PARPi sensitivity by preventing DNA end resection, thereby limiting HR repair. However, these same factors protect against IR-induced damage by facilitating efficient non-homologous end joining (NHEJ) repair of DSBs (*27*). Similarly, poly(ADP-ribose) glycohydrolase (PARG) promotes PARPi sensitivity by depleting cellular PAR chains, but its presence shields cells from IR-induced DNA damage (*28*). These observations highlight the complexity of therapeutic responses, where factors conferring sensitivity to one treatment may paradoxically drive resistance to another. Apart from restoration of BRCA1/2 function through genetic reversion, the molecular mechanisms that simultaneously contribute to both PARPi and IR sensitivity remain largely unexplored. One such mechanism might involve the promotion of RF degradation, a hallmark of therapy sensitivity of HRD cancers.

In this context, our study identifies Thousand And One Amino Acid Kinase 1 (TAOK1) as a novel factor that drives sensitivity to both PARPi and IR by regulating RF stability. TAOK1 is a MAP-kinase family member well described for its role in regulating Hippo-signaling and tissue growth (*29*, *30*). Recent findings by Zhu et al. (*31*) suggested that TAOK1 facilitates HR and that its inhibition with the ATP-competitive small molecule inhibitor CP43 enhances PARP sensitivity. Mechanistically, we show that TAOK1 interacts with PCNA and regulates ISG15 protein levels which are both known players in RF stability. TAOK1 aggravates RF degradation and replication gap formation and consequently triggers chemo- and radio-sensitivity.

## Results

### TAOK1 is critical for PARPi and radiotherapy sensitivity in BRCA1/2-deficient cells

To identify genes that modulate sensitivity to both PARPi and IR in BRCA-deficient cells, we carried out two complementary loss-of-function genetic screens. A kinome-wide shRNA screen targeting 652 kinases was performed in BRCA2;p53-deficient (*K14cre;Brca2^-/-^;p53^-/-^;* KB2P) mouse mammary tumor cells. Following transduction of the shRNA library, cells were treated with olaparib at an inhibitory concentration of 90 percent and after 10 days of treatment, genomic DNA of surviving cells was isolated and subjected to deep sequencing (Fig. 1A). In parallel, we performed additional screens applying RT as selective pressure. For this purpose, whole genome CRISPR-Cas9 screens were performed in RPE1-hTERT *BRCA1^-/-^;p53^-/-^*cells using the Brunello library (*32*). Successfully transduced cells were selected with a low dose of 1 or 2 Gy of IR over the course of three weeks (Fig. 1A). Sequencing data from the kinome and whole genome screens were analyzed with the MAGeCK algorithm (*33*) and both screens yielded high-quality results: as expected, loss of the master regulating kinase ATR emerged as a sensitizer to PARPi in BRCA2-deficient cells (*34*) (Fig. 1B and data S1). Moreover, key non-homologous end-joining (NHEJ) genes (*e.g. PRKDC, LIG4, NHEJ1, SHLD1, SHLD2, XRCC4*) were depleted under IR treatment, validating screen robustness (Fig. 1C, fig. S1A and data S2) (*35*). We thus concluded functionality of the screen to identify factors whose loss is causing therapy resistance. Notably, Thousand And One Amino Acid Kinase 1 (TAOK1) emerged as a shared hit across screens, and its depletion appeared to confer resistance to both olaparib and IR (Fig. 1B and C). We validated our findings by creating CRISPR-Cas9-mediated *Taok1* knockouts (KO) in BRCA1- and BRCA2-deficient mouse mammary tumor cells (KB1P-G3, KB2P-3.4 and KB2P-1.21) as well as in the human BRCA1-deficient MDA-MB-436 cancer cells (*36*). The KO efficacy was assessed by TIDE analysis and Western blotting (*37*), which confirmed complete loss of the TAOK1 protein in the KB1P-G3, KB2P-3.4 and MDA-MB-436 monoclonal *TAOK1* KO cells (fig. S1B, C and D). Partial protein reduction and frameshift modifications were detected in the polyclonal KB2P-1.21 cells (fig. S1E and F). Upon loss of TAOK1, all models exhibited resistance to both olaparib and IR, as shown by long term growth assays (Fig. 1D-F and fig. S1G-J). The reduced resistance levels in the polyclonal KB2P-1.21 cells likely reflected residual wild-type (wt) and in-frame alleles (fig. S1K, L). Importantly, re-introduction of HA-tagged *Taok1* cDNA (pOZ-Taok1) restored TAOK1 protein expression (fig. S1C) and re-sensitized KB1P-G3 cells to both PARPi and IR (Fig. 1E, F and fig. S1G, H), confirming the TAOK1 specificity of the resistance phenotype. Next, we aimed to validate the loss of *Taok1* and PARPi response *in vivo.* We therefore transduced BRCA1-deficient 3D mouse mammary organoids (KB1P-4S) with either NT or *Taok1* sgRNA and confirmed the modification rates on the *Taok1* gene using TIDE analysis and immunohistochemistry (fig. S1M, N). As shown in fig. S1O, loss of *Taok1* provided a strong growth advantage to KB1P-4S organoids treated with PARPi. To validate our findings *in vivo*, the transduced organoids were then orthotopically transplanted into mice. Daily treatment of either vehicle or olaparib (100mg/kg) for 56 consecutive days was initiated on mice bearing established tumors (50–100 mm^3^). Notably, *Taok1* depletion led to faster regrowth of tumors and resulted in accelerated mammary tumor-related morbidity (Fig. 1G). Taken together, these data establish TAOK1 as a critical modulator of PARPi and IR sensitivity in BRCA1/2-deficient cells.

**Fig. 1.**
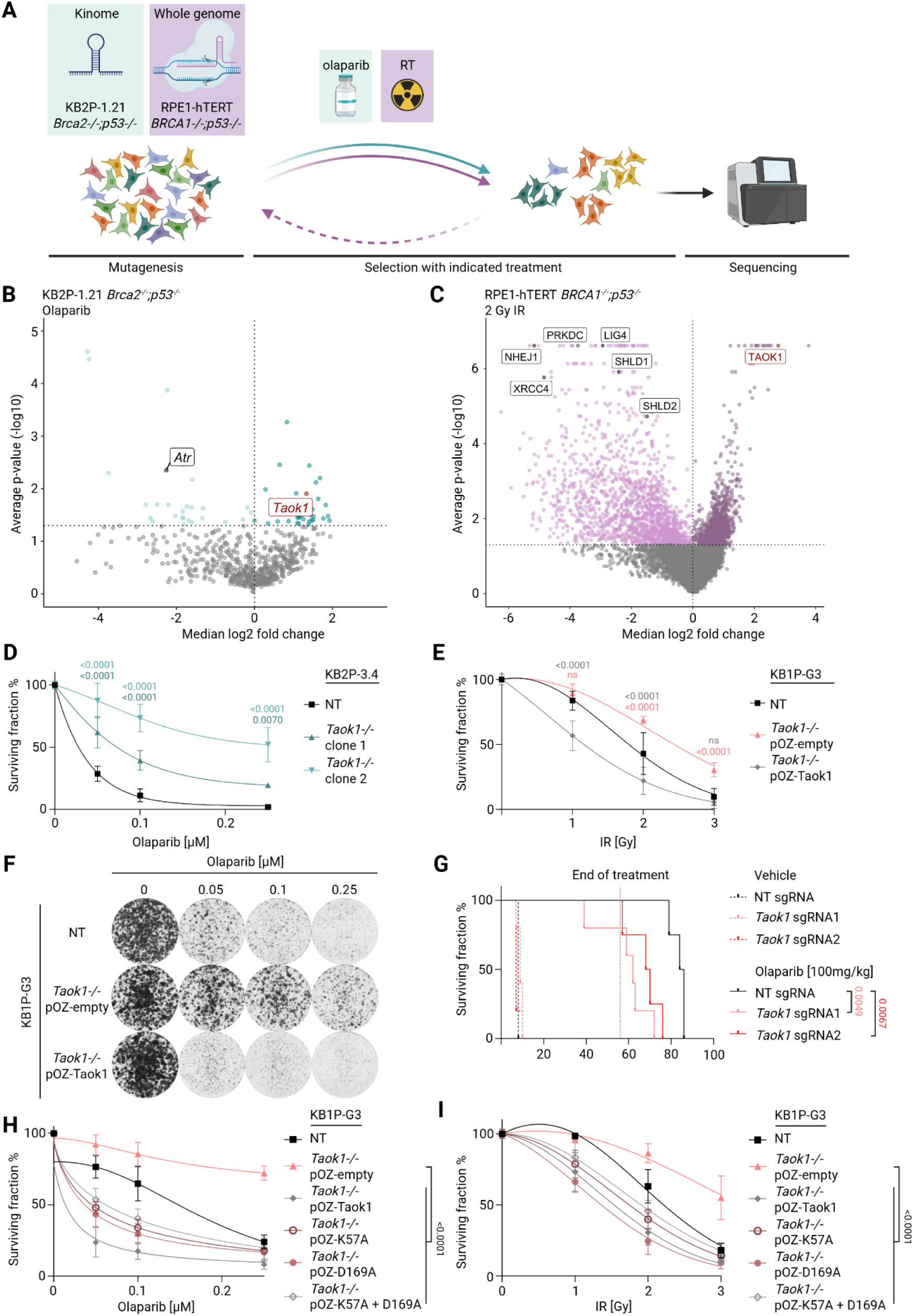
Functional genetic screens reveal loss of *Taok1* conferring PARPi and RT resistance. (**A**) Schematic outline of the kinome shRNA and whole genome CRISPR-Cas9 screens. (**B**) Volcano plot representing gene summary of MAGeCK analysis of the kinome shRNA screens in KB2P-1.21 cells treated with olaparib. (**C**) Volcano plot representing gene summary of MAGeCK analysis of the whole genome CRISPR-Cas9 screens in RPE1-hTERT *BRCA1^-/-^;p53^-/-^*cells treated with 2 Gy IR. (**D, E**) Growth assays in KB2P-3.4 (D) or KB1P-G3 (E) *Taok1^-/-^* cell lines treated with the indicated doses of olaparib (D) or IR (E). Statistical analysis was performed using two-way ANOVA followed by Dunnett’s test. (**F**) Representative images of growth assays in KB1P-G3 NT, *Taok1^-/-^* with empty rescue construct (pOZ-empty) or *Taok1* full cDNA rescue construct (pOZ-Taok1) treated with the indicated doses of olaparib. (**G**) Kaplan-Meier survival curve of mice transplanted with KB1P-4S organoids with NT or *Taok1* sgRNAs and treated with olaparib. Statistical analysis was performed with the log-rank test. (**H, I**) Growth assays in KB1P-G3 *Taok1^-/-^* cell lines complemented with indicated rescue construct. Cells were treated with indicated doses of olaparib (H) or IR (I). Statistical analysis was performed using two-way ANOVA followed by Dunnett’s test.

### TAOK1 drives therapy sensitivity independent of its kinase activity

To assess the role of TAOK1 kinase activity, we employed CP43, a selective inhibitor of this kinase. We tested a range of CP43 concentrations in combination with olaparib in our mouse mammary tumor models. Consistent with the findings of Zhu et al. (*31*), we observed a mild sensitivity in both BRCA1-proficient KB1P-G3B1 and BRCA1-deficient KB1P-G3 cells upon CP43 treatment (fig. S2A, B, C). Interestingly, the observed sensitization was also present in the KB1P-G3 *Taok1* KO cells (fig. S2B, C). CP43 is known to also inhibit TAOK2, a closely related paralogue of TAOK1, which might explain the sensitivity in the *Taok1* KO cell lines and the differential response compared to the CRISPR KO cell lines treated with olaparib. Additionally, the requirement for high doses of CP43 to elicit effects across all models may reflect nonspecific cellular toxicity. Together, these data show that the inhibition of kinase activity of TAOK1 using CP43 in BRCA1-deficient backgrounds does not reflect genetic *Taok1* depletion. An explanation for this result may be that the kinase function of TAOK1 is not responsible for the role of TAOK1 in conferring sensitivity of BRCA1/2-deficient cells to PARPi and IR.

To directly test whether kinase activity of TAOK1 is required for its role in sensitizing BRCA-deficient cells to DNA damage, we generated kinase-dead TAOK1 mutants. More specifically, we replaced the lysine at position 57 and/or the aspartic acid at position 169 with alanine (K57A, D169A), as they were previously described to abolish TAOK1 kinase activity (*38*, *39*). We re-introduced the K57A, D169A and the double mutant TAOK1 constructs in KB1P-G3 *Taok1* KO cells (fig. S2D) and tested the kinase activity by performing *in vitro* kinase assays upon pull-down of either WT (pOZ-Taok1) or kinase mutants (pOZ-K57A, pOZ-D169A, pOZ-K57A+D169A). As expected, we observed a reduction of kinase activity in all mutant constructs (fig. S2E). However, complementation of TAOK1 KO cells with the kinase mutants was still able to rescue sensitivity to both PARPi and IR, even at a low protein expression level (Fig 1H, I and fig. S2F, G). This demonstrates that the underlying mechanism of resistance in TAOK1 KO cells is not dependent on the kinase activity of TAOK1. Instead, TAOK1 likely acts through a kinase-independent scaffolding mechanism, perhaps by organizing protein complexes at sites of DNA damage.

### Loss of TAOK1 does not restore HR in BRCA1-deficient cells

To investigate whether TAOK1 loss influences PARPi and IR sensitivity through modulation of genomic instability, we assessed micronucleus formation as a surrogate marker. Notably, *Taok1* KO cells exhibited reduced micronucleus formation following DNA damage, indicating lower levels of genomic instability in these cells. (Fig. 2A). Because HR restoration can reduce micronuclei formation by promoting faithful repair of DSBs and limiting chromosomal mis-segregation, and since PARPi resistance frequently involves HR restoration, we investigated whether loss of TAOK1 promotes HR in BRCA-deficient cells. We therefore assessed RAD51 foci formation as a surrogate readout of HR proficiency in BRCA1/2-deficient cells upon both RT and PARPi treatment. In contrast to BRCA2-deficient cells, in BRCA1-deficient *Taok1^-/-^* cells RAD51 foci were increased after treatment with olaparib or IR (Fig. 2B, C and fig. S3A). However, this did not correlate with HR restoration. *BRCA1* or *BRCA2* knockdown strongly suppressed HR activity measured with a DSB-reporter (*40*), while co-depletion with *TAOK1* did not rescue this defect (Fig 2D and fig. S3B).

**Fig. 2.**
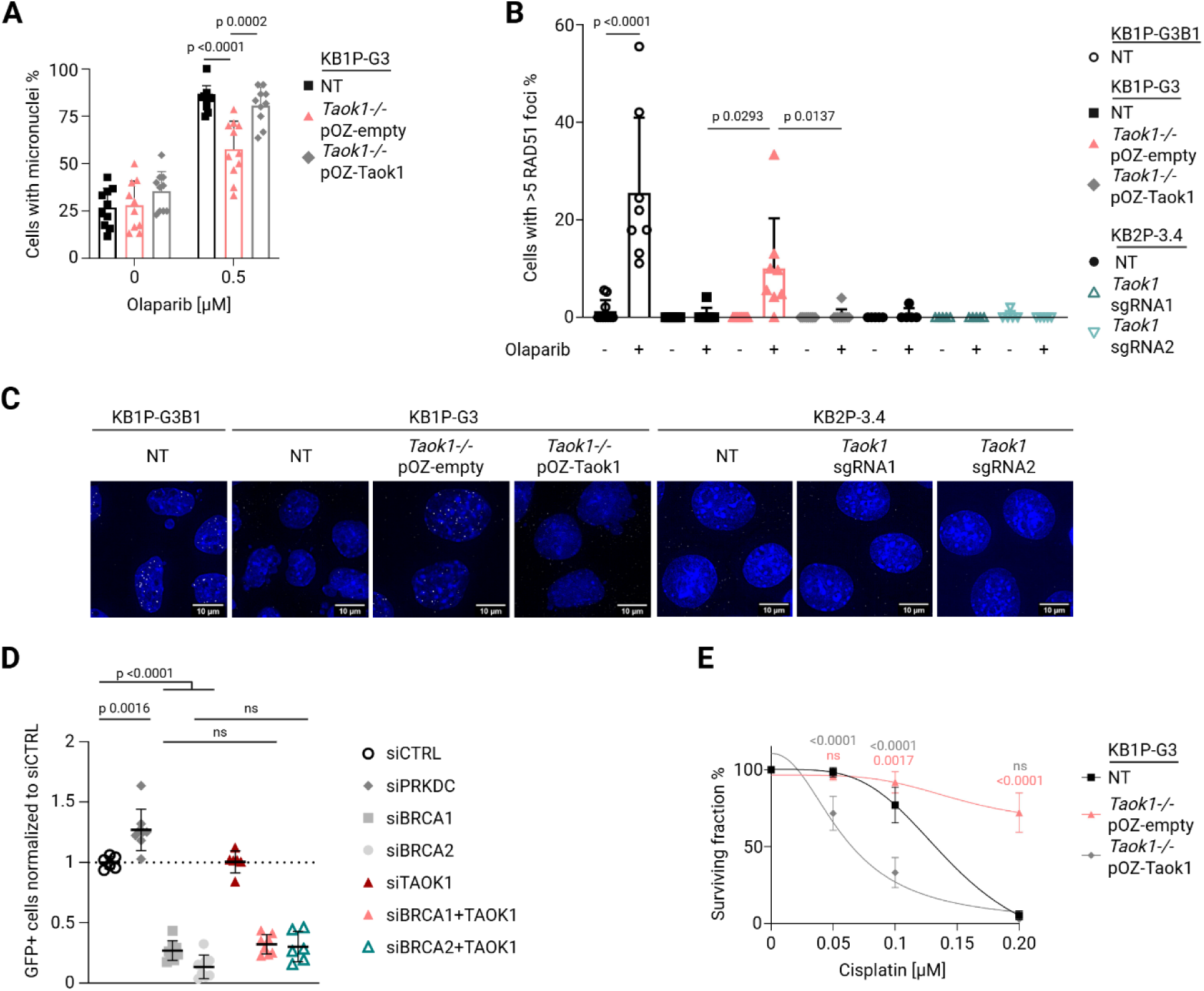
Loss of TAOK1 does not promote HR restoration in BRCA1-deficient cells. (**A**) Graph representing percentage of cells with micronuclei upon 0.5 µM olaparib treatment for 48 h. Bars are plotted as mean ± SD, statistical analysis was performed using ANOVA followed by Tukey’s multiple comparison test. (**B**) Quantification of RAD51 foci formation upon 10 µM olaparib. Bars represent mean ± SD, statistical analysis was performed using ANOVA followed by Tukey’s multiple comparison test. (**C**) Representative images of RAD51 foci formation upon 10 µM olaparib in *Brca1* reconstituted KB1P-G3B1, KB1P-G3 and KB2P-3.4 cell lines with the indicated modifications. (**D**) Quantification of DSB spectrum assay in HEK293T cells treated with the indicated siRNAs. Each dot represents an experiment ran in duplicates, bars represent mean ± SD, statistical analysis was performed using ANOVA followed by Tukey’s multiple comparison test. (**E**) Growth assays of KB1P-G3 NT, *Taok1^-/-^* and TAOK1 rescue cells treated with the indicated doses of cisplatin. Statistical analysis was performed using two-way ANOVA followed by Dunnett’s test.

These results confirm that TAOK1 loss does not restore canonical HR in BRCA1/2-deficient cells. Moreover, the resistance profile contrasts with that of PARPi-resistant cells arising from loss of the 53BP1-RIF1-REV7-shieldin-CST pathway, which become more radiosensitive (*27*). Instead, *Taok1* depletion conferred resistance not only to PARPi and IR, but also to platinum-based drugs such as cisplatin and carboplatin, which primarily cause replication-associated DNA damage (Fig. 2E, fig. S3C-E).

Taken together, these data indicate that the observed resistance in TAOK1- and BRCA1/2-deficient cells does not arise from HR restoration, but rather from altered responses to replication-associated stress. The RAD51 foci detected in BRCA1;p53-deficient *Taok1^-/-^*cells may therefore represent RAD51 functions at replication forks (*41*, *42*), which do not become easily visible in BRCA2-deficient cells.

### TAOK1 loss restores RF stability via PCNA and regulation of ISG15 expression

When we probed clinical tumor samples of cancer patients for TAOK1 expression by immunohistochemistry, we frequently observed a nuclear localization in addition to the cytoplasmatic localization (fig. S4A, B). Furthermore, TAOK1 was detected in protein lysates of both cytoplasmatic as well as nuclear fraction of KB1P-G3 cells (fig. S4C). In the absence of antibodies that detect mouse TAOK1 by immunofluorescence, the HA-tagged TAOK1 allowed us to look at its cellular localization in our mouse mammary tumor cells. Also in these cells, we observed that TAOK1 is present both in the cytoplasm, and in the nucleus, suggesting a separate function in addition to its role as a cytoplasmatic MAPK (fig. S4D).

Based on the nuclear localization of TAOK1 and the occurrence of RAD51 foci in BRCA1-deficient *Taok1^-/-^* cells, we hypothesized that TAOK1 might play a regulatory role at replication forks. We therefore interrogated whether TAOK1 is localized to DNA replication sites by performing an *in situ* analysis of protein interactions at DNA replication forks (SIRF) (*43*). Indeed, HA-TAOK1 robustly co-localized with EdU-labeled replication sites, both under basal and hydroxyurea-induced replication stress conditions (Fig. 3A, B), indicating fork interactions independent of acute damage and a role of TAOK1 in basal replication dynamics.

**Fig. 3.**
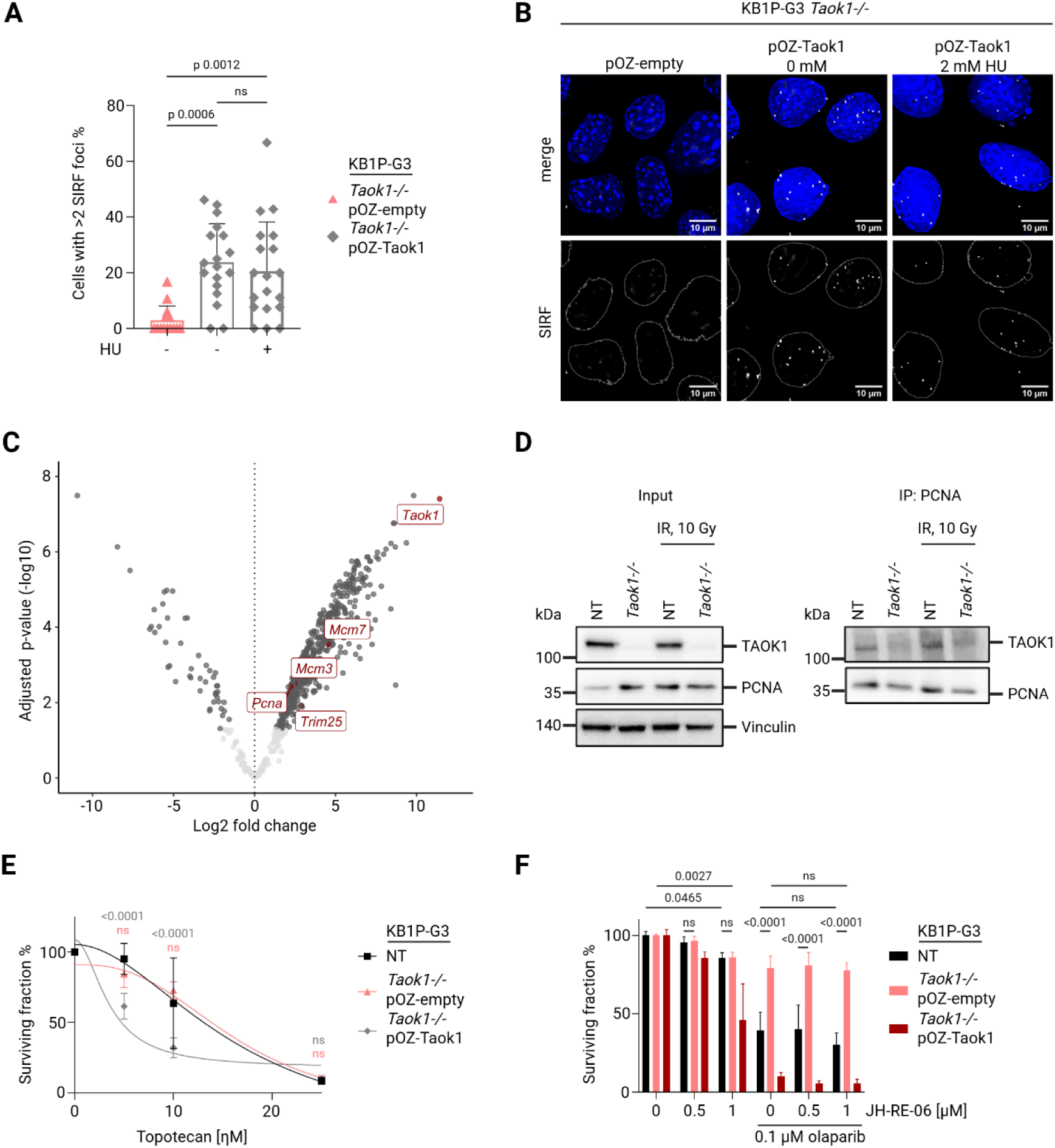
TAOK1 is present at replication sites and interacts with PCNA. (**A**) Quantification of SIRF foci formation of HA-tagged TAOK1 with and without 2 mM HU. Bars represent mean ± SD, statistical analysis was performed using ANOVA followed by Tukey’s multiple comparison test. (**B**) Representative images of analysis shown in (**A**). (**C**) Volcano plot of mass spectrum analysis of TAOK1 co-IP samples from KB1P-G3 NT nuclear fraction compared to TAOK1 KO. Positive log2 fold change values indicate enrichment in the NT nuclear fraction compared to TAOK1 KO of four independent replicas. (**D**) Western blot of input and co-IP of KB1P-G3 NT and TAOK1 KO. Pulldown was performed with PCNA antibody. (**E**) Growth assay of KB1P-G3 NT, TAOK1 KO and TAOK1 rescue cells treated with topotecan at indicated doses. Statistical analysis was performed using two-way ANOVA followed by Dunnett’s test. (**F**) Growth assay of KB1P-G3 NT, TAOK1 KO and TAOK1 rescue cells treated with JH-RE-06 alone or in combination with olaparib. Statistical analysis was performed using two-way ANOVA followed by Tukey’s test.

While the cytoplasmatic role of TAOK1 as MAPK has been already extensively studied (*29*, *30*, *44*), its nuclear function is unclear. To understand how TAOK1 influences RF dynamics and therapy sensitivity, we sought to identify its nuclear binding partners. To this purpose, we co-immunoprecipitated (co-IP) the nuclear fraction of cells expressing wildtype TAOK1 and whole cell lysates of *Taok1^-/-^* cells as negative control using magnetic beads coated with a TAOK1 antibody and subsequently digested and analyzed the purified proteins by mass spectrometry (Data S3). The volcano plot depicted in Fig. 3C shows the significant interaction partners of four independent co-IP experiments. Among these, TAOK1 was one of the significant hits, which served as a positive control proving functional pulldown (Fig. 3C). Based on our previous observations, we examined the results specifically for replication-associated proteins. Among the significantly enriched interactors, we identified the replication clamp PCNA and the helicase subunits MCM3 and MCM7, components of the CMG complex, highlighting a potential role for TAOK1 in replisome regulation (Fig. 3C). PCNA as interaction partner is especially interesting as its presence at replication sites is regulating downstream pathways responsible for DDT and RF protection, thereby influencing PARPi response particularly in BRCA-deficient cells (*16*, *17*, *19*). We validated the interaction between TAOK1 and PCNA by co-IP and immunoblotting (Fig. 3D). Notably, TAOK1-deficient cells showed no resistance to topoisomerase I inhibitors (topotecan, camptothecin), which do not promote PCNA ubiquitination (*45*), suggesting that TAOK1 specifically affects PCNA-regulated DNA damage responses (Fig. 3E and fig. S4E).

Ubiquitinated PCNA (ub-PCNA) is important for the induction of DDT pathways by promoting TLS and TS (*16*, *17*). Moreover, activation of DDT pathways is increased in tumors and associated with drug resistance (*46*, *47*). Activation of the TS pathway was excluded as potential mechanism of resistance in TAOK1-deficient cells as TS relies on BRCA1/2 for loading of RAD51 filaments (*48*, *49*). To investigate whether loss of *Taok1* is specifically triggering TLS, we explored the response of *Taok1^-/-^*cells to the TLS inhibitor JH-RE-06. Using growth assays in which cells were treated with the inhibitor alone or in combination with olaparib did not show major differences of response, suggesting that activation of TLS is not the major pathway of olaparib resistance in BRCA1-deficient TAOK1 KO cells (Fig 3F). The same experimental setup using BRCA2-deficient KB2P-3.4 cells demonstrated that TLS in KB2P-3.4 NT is more important for cell survival while KB2P-3.4 *Taok1^-/-^* cells hardly showed any sensitivity to JH-RE-06 alone or in combination with olaparib (fig. S4F).

Based on the role of PCNA in protecting RF and its interaction with TAOK1, we examined RF dynamics in TAOK1 KO cell lines. When we investigated replication speed of untreated NT versus *Taok1^-/-^* cells by measuring total CldU/IdU track length, we did not observe any difference (Fig. 4A). However, loss of *Taok1* rescued the susceptibility of BRCA1/2-deficient cells (*10*) to fork degradation upon HU-induced DNA damage to a similar extent as BRCA1-expressing cells, suggesting that loss of *Taok1* protects replication forks from nucleolytic degradation (Fig. 4B and fig. S4G). In light of the emerging role of replication gaps in PARPi vulnerability and DDT pathways (*50*), we also analyzed the accumulation of ssDNA gaps in *Taok1^-/-^* cells. For this purpose, we treated KB1P-G3 NT and *Taok1^-/-^* cells with olaparib 2h prior to and during labeling with CldU and IdU and added S1 nuclease to digest ssDNA gaps, which can be detected by shortening of the IdU tract in BRCA1-deficient cells. As depicted in fig. S4H, BRCA1-deficient *Taok1^-/-^* cells accumulated fewer ssDNA gaps as shown by the unabridged length of IdU tract upon S1 nuclease treatment (Fig. 4C). Consistently, human BRCA1-deficient *Taok1⁻/⁻*MDA-MB-436 cancer cells also displayed reduced ssDNA gap accumulation under the same conditions (Fig. 4C). Complementary assays using RPA foci quantification (Fig. 4D, E and fig. S5A, B) and native BrdU incorporation (fig. S5C, D) also demonstrated reduced ssDNA formation in BRCA1/2-deficient cells.

**Fig. 4.**
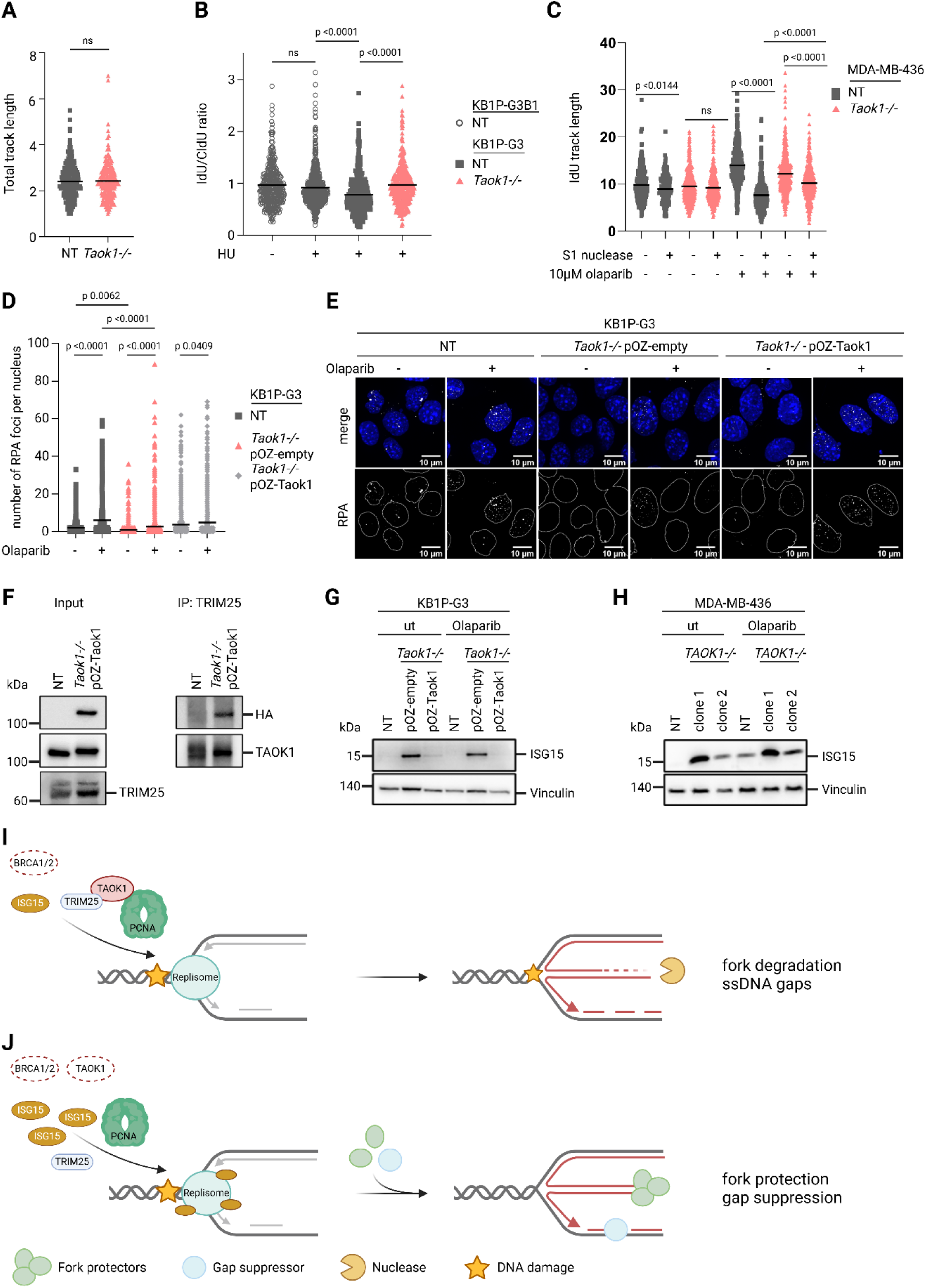
TAOK1 loss restores RF stability. (**A**) Graph showing replication speed by total track length of untreated DNA fiber assay in KB1P-G3 NT and TAOK1 KO cells. Bar represents the mean of at least 250 fibers, statistical analysis was done with unpaired t-test. (**B**) DNA fiber analysis in indicated cell lines with or without 4 mM HU treatment. Bar represents the mean of IdU/CldU ratio of at least 300 fibers, statistical analysis was done with unpaired t-test. (**C**) DNA fiber analysis in indicated cell lines treated with 10 µM olaparib 2h prior and while labelling with CldU and IdU, ssDNA was digested with S1 nuclease if indicated. Bar represents the mean of IdU track length of at least 250 fibers, statistical analysis was done with ANOVA followed by Tukey’s multiple comparison test. (**D, E**) Quantification (D) and representative images (E) of immunofluorescence analysis of RPA foci formation in KB1P-G3 NT, TAOK1 KO and TAOK1 rescue cells upon olaparib treatment (10 µM for 16 h). Bars indicate the mean number of foci per nucleus of at least 1000 nuclei. Statistical analysis was done with ANOVA followed by Tukey’s multiple comparison test. **(F)** Western blot of input and co-IP of KB1P-G3 NT and TAOK1 rescue cells. Pulldown was performed with TRIM25 antibody. (**G, H**) Western blot of indicated KB1P-G3 (H) and MDA-MB-436 (I) cell lines for ISG15 protein levels. Cells were untreated or treated with 10 µM olaparib for 24 h. Vinculin was used as loading control. (**I, J**) Schematic predicting replication dynamics in presence (I) or absence of TAOK1 (J) in BRCA1/2-deficient cells. (I) TAOK1 interacts with PCNA and TRIM25 and does not allow alternative fork protection mechanisms or ssDNA gap repair in BRCA-deficient cells, leading to RF instability. (J) In the absence of TAOK1, increased ISG15 expression and potentially recruitment to RF activates fork protection and gap suppression mechanisms and thus, RF stability.

In addition to ubiquitination of PCNA, PTMs by ISG15 conjugation have been shown to regulate downstream tolerance pathways in a similar manner. Notably, ISGylation of PCNA has been reported to limit TLS activity (*20*). Beyond its interaction with PCNA, ISG15 appears to target multiple proteins at replication sites, as it has been implicated in preserving RF integrity through the ISGylation of factors at nascent DNA (*21*, *51*). Although we did not observe a significant enrichment of ISG15 itself in the pull-down of TAOK1, we detected TRIM25, a ubiquitin E3-ligase that also acts as ISG15-ligase (Fig. 3C) (*52*). Indeed, we confirmed the interaction of TAOK1 with TRIM25 by co-IP and

PLA (Fig. 4F and fig. S5E). Interestingly, we also observed an increase of free ISG15 in *Taok1^-/-^* cells compared to NT control cells (Fig. 3G, H) in untreated conditions as well as after induction of replication stress by olaparib, suggesting a potential role for TAOK1 in modulating ISG15 conjugation dynamics. Overall, these findings suggest a role for TAOK1 as regulator of the replisome. While PCNA ubiquitination does not seem to be crucial in TAOK1 KO cells, we hypothesize that regulation of ISG15 on PCNA and potentially other proteins on nascent DNA might influence replication fork integrity.

Together, these data reveal that TAOK1 localizes to RF and promotes RF degradation and gap formation in BRCA1/2-deficient cells. Loss of TAOK1 stabilizes forks potentially by increased ISGylation of the replisome and mitigates replication-associated DNA damage, contributing to resistance to therapies targeting fork vulnerability (Fig. 4I, J).

### High TAOK1 expression levels correlate with chemotherapy sensitivity

The functionality of the TAOK1 antibody for immunohistochemistry in patient samples (fig. S4A), along with the observed variation in staining intensity across different tumor types (fig. S4B), suggests that TAOK1 expression could serve as a predictive biomarker for therapy response. We therefore, consulted the Swedish Cancerome Analysis Network – Breast (SCAN-B) dataset for breast cancer patients (*53*, *54*). Overall survival (OS) of TAOK1 high versus TAOK1 low expression in the whole cohort revealed no significant difference (Fig. 5A). As shown in Fig. 1D and fig. S1G-I, K, O, S3C-E, TAOK1 drives response to several DNA damage-inducing agents, including PARPi, platinum drugs, in addition to IR. Since DNA damage-inducing chemotherapy is frequently used to treat breast cancer patients, we therefore hypothesized that TAOK1 expression levels predict treatment outcome of patients treated with chemotherapy specifically. Indeed, further evaluation of the dataset with patients treated with chemotherapy showed a significant survival advantage for patients with high TAOK1 expression (Fig. 5B), while TAOK1 expression was not associated with response in patients who did not receive chemotherapy treatment (fig. S5F). Therefore, our findings indicate that evaluating TAOK1 expression in patients could serve as a useful predictor of response to DNA-damage-inducing chemotherapy.

**Fig. 5.**
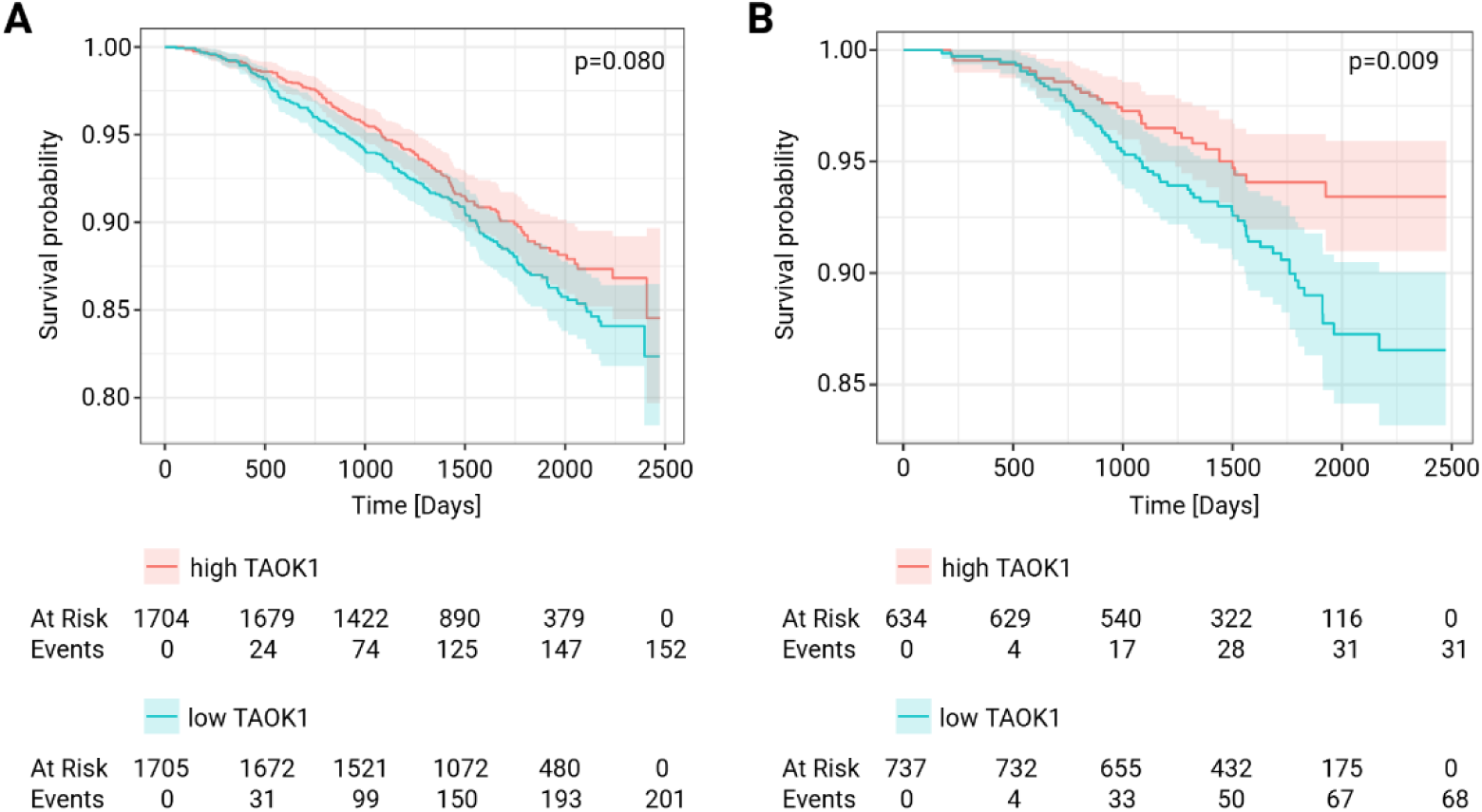
High TAOK1 expression levels correlate with chemotherapy response. (**A, B**) Analysis of the SCAN-B dataset for TAOK1 high or low expression. Kaplan-Meier curves for survival probability of all patients (A) and patients treated with chemotherapy specifically (B). Risk tables are shown below. Statistical analysis was done with log-rank test.

## Discussion

Our study identifies TAOK1 as a novel regulator of RF dynamics that modulates sensitivity to DNA damage-inducing treatments in BRCA1/2-deficient cells. While previous work to understand therapy sensitivity has focused on established DNA repair proteins that govern HR or NHEJ, our findings uncover a distinct, kinase-independent role for TAOK1 in replication-associated stress response. Specifically, we show that TAOK1 promotes RF degradation and replication gap formation, thereby enhancing the cytotoxic effects of PARPi, IR, and platinum-based agents.

Unlike factors such as the 53BP1-RIF1-REV7-shieldin-CST pathway, which mediate PARPi sensitivity by blocking DNA end resection but protect against IR by facilitating NHEJ (*22*, *24–26*, *55*, *56*), TAOK1 promotes sensitivity to both PARPi and IR. Importantly, TAOK1 depletion conferred resistance to multiple genotoxic therapies without restoring HR capacity, distinguishing it mechanistically from many other modulators of therapy resistance. Although RAD51 foci were detected in TAOK1-deficient *Brca1^-/-^;p53^-/-^*cells, functional assays confirmed that canonical HR was not restored, and resistance extended to platinum drugs, which primarily induce replication-associated damage. We hypothesize that these RAD51 foci are related to RAD51 function at RF, for which, similarly to double strand breaks, the presence of BRCA2 is indispensable for their loading (*57*). Consistent with this, our findings demonstrate that TAOK1 influences RF stability by promoting the formation of ssDNA gaps and facilitating RF degradation, mechanisms that heighten the vulnerability of BRCA1/2-deficient cells to genotoxic stress (*50*).

Mechanistically, we demonstrate that TAOK1 localizes to RF and interacts with PCNA and the ISG15-ligase TRIM25. The interaction of TAOK1 and PCNA further implicates it in the regulation of post-replicative DDT pathways. While TAOK1 does not appear to hyperactivate TLS, its potential role as regulator of ISGylation of PCNA and other proteins on nascent DNA supports a function in replication fork integrity. This is further evidenced by the fact that TAOK1 leads to RF degradation and suppresses replication gap repair.

In line with a recent study by Zhu *et al.*, which proposed that TAOK1 supports HR and that TAO kinase inhibition could enhance PARPi response (*31*), we observed mild sensitivity to the ATP-competitive TAOK inhibitor CP43 in both wild-type and BRCA1;p53-deficient cells. However, the inhibitor does not exactly mimic genetic depletion of TAOK1, as TAOK1 KO BRCA1;p53-deficient cells were still responding to CP43. Although these discrepancies may reflect distinct biological contexts, they may also be explained by our data showing that the kinase activity of TAOK1 is not mediating the DNA damage-induced drug sensitivity. We provide genetic evidence that kinase-dead TAOK1 mutants fully restore therapy sensitivity, strongly supporting a kinase-independent scaffolding role.

TAOK1 is involved in various biological processes associated with Hippo signaling, including cytoskeleton stability (*38*), neuronal development (*58*), chromosome segregation during mitosis (*59*, *61*) and cancer progression (*29*, *30*, *60*). Here, we expand its repertoire to include replication stress responses in the nucleus. Intriguingly, TAOK1 loss reduced micronuclei formation, an indicator of chromosomal instability, despite previous reports linking TAOK1 to mitotic chromosome segregation errors (*59*, *61*). This paradox can be explained by the HR deficiency context in which we studied TAOK1, and may reflect the complex crosstalk between replication stress, fork stability, and chromosomal maintenance in BRCA-deficient settings.

Regarding therapeutic implications, our findings are twofold. First, TAOK1 could serve as a biomarker for response to PARPi, RT, and platinum therapies in HR-deficient tumors. In particular, as shown by the SCAN-B dataset, TAOK1 expression levels might predict chemotherapy response. Second, strategies aimed at upregulating or stabilizing TAOK1, rather than inhibiting it, may sensitize resistant BRCA1/2-deficient tumors to DNA-damaging agents. However, the lack of specificity of current TAOK1-targeting drugs, and the kinase-independent nature of its function, highlight the need for novel therapeutic strategies that may enhance scaffolding interactions of TAOK1, particularly with PCNA.

In summary, this study identifies TAOK1 as a non-catalytic modulator of replication fork dynamics and therapy sensitivity in BRCA1/2-deficient cancers. Further elucidation of its protein interactions and structural domains may pave the way for new strategies to exploit replication stress in precision oncology.

## Materials and Methods

### Cell culturing conditions

The KB1P-G3 cell line was established from *K14cre;Brca1^F/F^;p53^F/F^*mouse mammary tumors as previously described (*22*). KB1P-G3 cell line was reconstituted with human BRCA1 cDNA as previously described generating the isogenic KB1P-G3B1 cell line (*27*). KB2P-3.4 and KB2P-1.21 cell lines were established from *K14cre;Brca2^F/F^;p53^F/F^* mouse mammary tumors as previously described (*63*). All these cell lines were cultured in Dulbecco’s Modified Eagle Medium/Nutrient Mixture F-12 (DMEM/F-12, Gibco) supplemented with 10% fetal calf serum (FCS, Sigma), 50 units/ml penicillin-streptomycin (Gibco), 5 μg/ml Insulin (Sigma,#I0516), 5 ng/ml cholera toxin (Sigma, #C8052) and 5 ng/ml murine epidermal growth-factor (EGF, Sigma, #E4127). All mouse mammary tumor cell lines were cultured at 3% O_2_. HEK293FT cell lines as well as Phoenix-ECO cell line were cultured in Dulbecco’s Modified Eagle Medium (DMEM, Gibco) supplemented with 10% fetal calf serum (FCS, Sigma) and 50 units/ml penicillin-streptomycin (Gibco). MDA-MB-436 cell line was grown in RPMI with L-Glutamine (Gibco) supplemented with 10% fetal calf serum (FCS, Sigma) and 50 units/ml penicillin-streptomycin (Gibco). RPE1-hTERT *BRCA1^-/-^;p53^-/-^*were grown at 3% O2 in DMEM (Gibco) supplemented with 10% fetal calf serum (FCS, Sigma), 50 units/ml penicillin-streptomycin (Gibco), 1x GlutaMAX (Gibco), 1x non-essential amino acids (NEAA; Gibco). KB1P-4S mouse mammary organoids were cultured in normal oxygen embedded in Culturex Reduced Growth Factor Basement Membrane Extract Type 2 (BME, Trevigen) in Advanced DMEM-F12 (Gibco) supplemented with 50 units/ml penicillin/streptomycin, HEPES 1M, 1x GlutaMAX (Gibco), N-acetylcysteine 125 µM (Sigma), B-27 (Gibco), 50 ng/ml murine epidermal growth-factor (EGF, Sigma, #E4127). All cell lines were regularly tested for mycoplasma contamination (Mycoalert, Lonza).

### In vivo studies

All animal experiments were approved by the Canton of Bern (Switzerland, license BE69/2021). KB1P-4S tumor organoid with NT or *Taok1* sgRNA modification were dissociated into single cells for transplantation, washed in PBS, resuspended in tumor organoid medium and mixed 1:1 with BME. Organoids suspension of a total of 10^5^ cells were injected in the fourth right mammary fat pad of 6-9 week-old female NMRI nude mice. Treatment of tumor bearing mice was initiated when tumors reached a size of 50-100 mm^3^. Tumor size was measured by caliper and volume was calculated as ((length x width^2^)/2). Mice were assigned into a vehicle-treated group (NT gRNA n=5, *Taok1* sgRNA1 n=5, *Taok1* sgRNA2 n=5) and olaparib-treated group (NT gRNA n=4, *Taok1* sgRNA1 n=5, *Taok1* sgRNA2 n=4). Olaparib was given intraperitoneally for 56 consecutive days at 100mg/kg. Animals were anesthetized with isoflurane, sacrificed with CO_2_ and cervical dislocation when tumors reached a volume of ∼1000 mm^3^. Maximum body weight loss permitted is 10% of the total animal weight.

### Drugs and reagents

The following chemical reagents were used throughout the study: olaparib (kindly provided by AstraZeneca and Syncom (Groningen, Netherlands)), talazoparib (Selleckchem, #S7048), cisplatin (Teva, #7680479980428), oxaliplatin (Teva, #7680616880017), carboplatin (Teva, Cat#6985451), camptothecin (Selleckchem, #S1288), topotecan (Selleckchem, #S9321), hydroxyurea (Sigma, #H8627), TAOK inhibitor CP 43 (MedChemExpress, #HY-112136), JH-RE-06 (TargetMol, # T15611) 5-Iodo-2’-deoxyuridine (IdU) (Sigma, #I7125), 5-Chloro-2’deoxyuridine (CldU) (Sigma, #C6891), Ethynyl-2’-deoxyuridine (EdU) (Sigma, #A10044).

### Antibodies

The following primary antibodies were used: rabbit anti-RAD51 (Bioacademia, #70-012, 1:1000), mouse anti-HA (Biolegend, clone 16B12, 1:1000), rabbit anti-HA (Cell Signaling #C29F4, 1:1000), mouse anti-Biotin (Jackson Immuno Research, #200-002-211, 1:200), mouse anti-BrdU/IdU (BD Biosciences, #347580, 1:50), rat anti-BrdU/CldU (Abcam, clone [BU1/75 (ICR1)], #ab6326, 1:250), rabbit anti-TAOK1 (Proteintech, #26250-1-AP, 1:1000), rabbit anti-Vinculin (Cell Signaling, #1390S, 1:1000), mouse anti-PCNA (Santa Cruz, #sc-56,-1:500), rabbit anti-PCNA (Cell Signaling, #13110S, 1:500), mouse anti-alpha tubulin (Sigma, #T9026, 1:1000), mouse anti-ISG15 (Santa Cruz, #sc-166755, 1:500), rabbit anti-BRCA1 (Cell Signaling, #9010, 1:1000), rabbit anti-BRCA2 (Cell Signaling, #10741, 1:1000), rabbit anti-DNA-PKcs (Cell Signaling, #4602, 1:1000), rabbit anti-H2AX (Cell Signaling, #2595, 1:1000), rabbit anti-TRIM25 (Proteintech, #12573-13AP, 1:1000).

### CRISPR-Cas9 screen

RPE1-hTERT *BRCA1^-/-^;p53^-/-^* cells were infected with genome-wide human lentiviral Brunello library with a transduction efficiency of 30% after selection with puromycin (as previously described (*64*), 20 µg/ml). The library contains 76,441 sgRNAs targeting 19,114 genes. 7 Days after puromycin selection 24 million cells were seeded to maintain a library coverage of 300x and Day 0 sample was collected. The experiment was performed in two technical replicates. Cells were either left untreated or treated with fractionated doses of IR (two doses of 1 or 2 Gy/week). Every 7 days, cells were trypsinized and 24x10^6^ cells were re-seeded and treated as week one. The experiment was collected after 3 weeks. Genomic DNA of surviving cells was isolated with MN Nucleo Bond XL Blood kit (Macherey-Nagel 740950.50) and submitted for Illumina sequencing at the Broad Institute (Cambridge, USA). Sequencing results were deconvolved into sgRNA abundance using PoolQ. Results from 2 independent replicas were analyzed with the MAGeCK (Model-based Analysis of Genome-wide CRISPR-Cas9 Knockout) algorithm (*33*).

### shRNA kinome screen

Kinome shRNA screen was performed as previously described (*28*). In brief, a custom shRNA library targeting 652 kinases with a total of 3255 hairpins was designed in pLKO.1 vector. The shRNA library was stably introduced into KB2P-1.21 cell line by lentiviral transduction. Cells were seeded at a coverage of 500x in 3 technical replicates and subsequently treated with olaparib at IC90. Cells were selected with olaparib for 10 days. Genomic DNA of surviving clones was extracted and sequenced. Analysis was performed using MAGeCK algorithm.

### Lentiviral transductions

Lentivirus was generated using HEK293FT cells. Therefore, 8x10^6^ cells were seed in 150 mm dishes and on the following day transfected with 3^rd^ generation lentiviral packaging plasmids and pLentiCRISPRv2 vector containing the sgRNA. The plasmid sequence was verified with Sanger sequencing. Transfection was performed by Calcium-precipitation using 2xHBS (280 nM NaCl, 100 mM HEPES, 1.5 mM Na_2_HPO_4_, pH 7.22), 2.5M CaCl_2_ and 0.1x TE-Buffer (10 mM Tris pH8.0, 1 mM EDTA pH8.0, diluted 1:10 with dH_2_O). The virus-containing supernatant was collected after 30 h and centrifuged with ultracentrifugation in a SW40 rotor at 20,000 rpm for 2 h. Virus pellet was resuspended in 100 µl PBS. The virus titer was determined using a qPCR Lentivirus Titration Kit (Applied Biological Materials, #LV900). Lentiviral CRISPR modification was performed in all KB1/2P mouse cell lines. Target cells for lentiviral transduction were seeded at 150,000 cells per well in a 6-well plate. On day 1, virus was added at multiplicity of infection (MOI) of 25 with 8 µg/ml polybrene (Merck). On day 2, virus containing medium was replaced with selection medium containing puromycin (3.5 µg/ml, Gibco) and cells were selected for 3 days. Tumor-derived organoids were transduced with lentivirus as previously described (*65*).

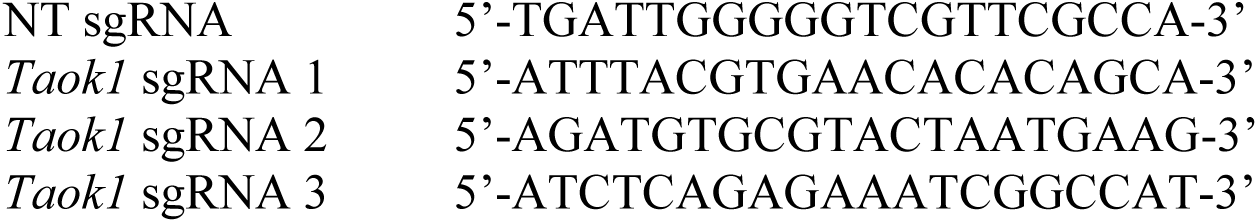

### Genome editing

For CRISPR-Cas9 editing of human MDA-MB-436 cell line, NT sgRNA and *TAOK1* sgRNAs were cloned into pX330 vector (Addgene, #42230) with puromycin resistance cassette. The plasmid sequence was verified with Sanger sequencing. Target cells were transfected using the TransIT-LT1 transfection reagent (Mirus, #MIR6604) following manufacturer’s protocol. Cells were selected with puromycin (1 µg/ml, Gibco) for 3 days.

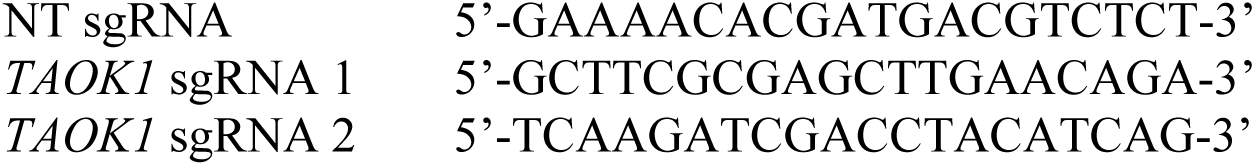

For the reconstitution of TAOK1 in KB1P-G3 *Taok1^-/-^* cell line, *Taok1* coding sequence of mus musculus was ordered from Eurofins and cloned into pOZ-N-FH backbone adding 1x HA tag at the N-terminus of *Taok1* sequence. Cloning was performed with in-fusion HD cloning kit (Takara, #12141). Site-directed mutagenesis for kinase mutant constructs pOZ-K57A, pOZ-D169A and pOZ-K57A+D169A was performed with GeneArt Site-Directed Mutagenesis kit (Invitrogen, #A13282) according to manufacturer’s protocol. Retrovirus was generated in HEK Phoenix ECO cell lines, pOZ -Taok1, pOZ -K57A, pOZ -D169A, pOZ-K57A+D169A or pOZ -empty plasmid were transfected using Turbofectin reagent (Origene, #TF81001). After 48 hours, supernatant containing retrovirus was collected and filtered through 0.45 µm filter. *Taok1^-/-^* KB1P-G3 target cells were seeded at 400,000 cells in a 10 cm dish and on day 1 retrovirus was added with 8 µg/ml polybrene (Merck). Selection of transduced cells expressing IL2Rα was performed on 70% confluent target cells with magnetic beads coated with CDC25 (Thermo Scientific, Dynabeads CDC25, #11157D).

### gDNA isolation, amplification, TIDE analysis

Modification rates of CRISPR-Cas9 edited cell lines were assessed by TIDE algorithm. Therefore, genomic DNA was isolated from cell lines using QIAmp DNA mini kit (Qiagen) according to manufacturer’s protocol. Target sequence was amplified with PCR using Phusion Flash High-Fidelity Master Mix (Thermo Scientific, #F-548L) according to manufacturer’s protocol. A 3-step PCR protocol was performed: 98 °C for 10’’, 30 cycles at 98°C for 1’’, 58°C for 5’’and 72°C for 15’’, final extension at 72°C for 1’. PCR product was purified with QIAquick PCR purification kit (Qiagen) following manufacturer’s protocol and submitted for Sanger sequencing with the corresponding forward primer. mus musculus

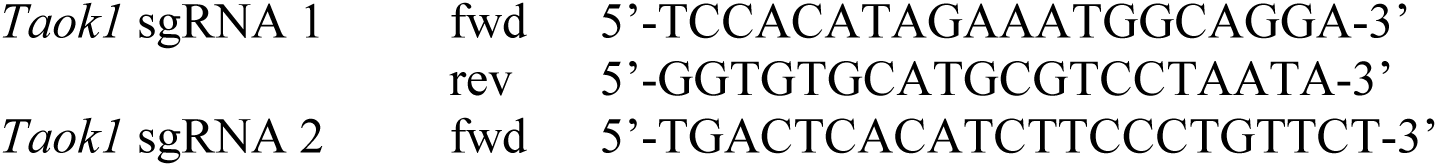
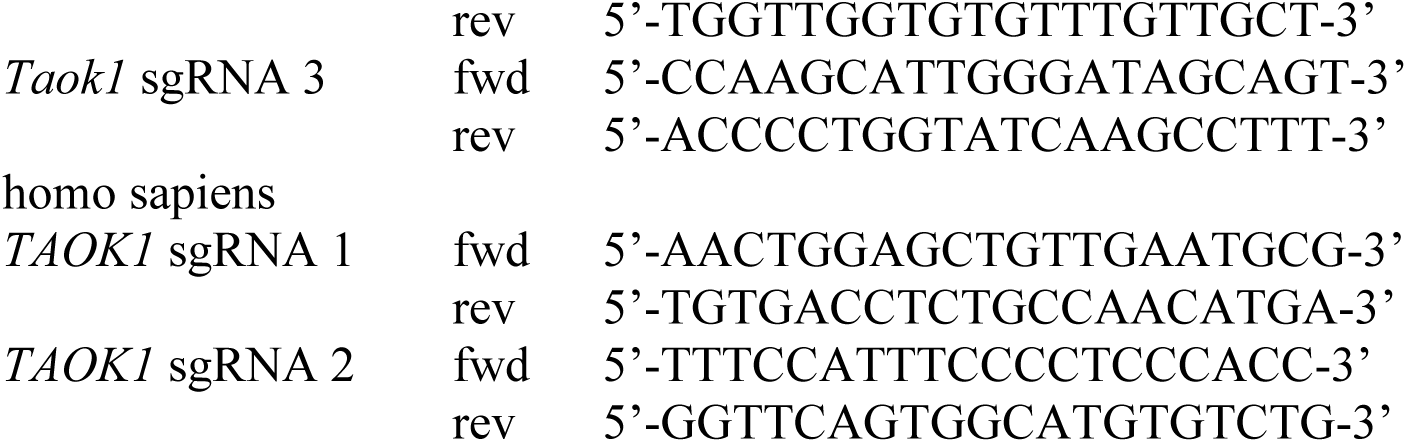

### Growth assays

To assess surviving fraction of cell lines treated with chemotoxic agents (olaparib, talazoparib, cisplatin, carboplatin, oxaliplatin, camptothecin) or IR in KB1/2P cell lines, cells were seeded in 6-well plates (4,000 cells/well). Cells were treated at indicated doses with the following schedule. PARPi and topotecan were added on day 1 after seeding and refreshed on day 4, platinum drugs were added on day 1 and removed after 24h, camptothecin was added on day 1 and left until fixation, all control group were treated with the same concentration of the respective drug diluent. IR was performed in a fractionated manner, cells were treated on three consecutive days after seeding with XRAD225 XL (Precision X-Ray) machine using Cupper filter. All plates were fixed on Day 8 with 4% para-formaldehyde and stained with 0.1% crystal violet. Surviving fraction was analyzed in an automated manner with ColonyArea plugin in ImageJ. Viability of MDA-MB-436 cell lines and topotecan treated KB1P-G3 cells was assessed in 24-well plate format. Cells were seeded at 800 cells/ well (KB1P-G3) or 2000 cells/well (MDA-MB-436). On day 8, cell viability was assessed after 4h incubation with CellTiter-Blue (Promega, #G8080). Fluorescence at was measured with Varioskan LUX (Thermo Scientific) plate reader.

### Immunoblotting

Cells were collected, washed with PBS and lysed in RIPA buffer (50 mM Tris/HCl pH 7.4; 1% NP-40; 0.5% Na-deoxycholate; 0.1% SDS; 150 mM NaCl, 2 nM EDTA, 50 mM NaF) with Halt protease inhibitor and phosphatase inhibitor cocktail (Thermo Scientific, #78420). Lysates were incubated for 30min on ice and vortexed every 5min. Samples were centrifuged at 14,000 rpm for 10min at 4°C and the supernatant was collected. Protein concentration was assessed with Pierce BCA protein Assay Kit (Thermo Scientific, #23225). Protein lysates were prepared with 6x SDS sample buffer and denatured at 95°C for 10min. Proteins were separated by 7.5% or 12% SDS/PAGE gel, transferred to 0.45 µm PVDF (Immobilon) or 0.22 µm nitrocellulose membrane (GE Healthcare) and blocked in 5% dry milk powder in TBS-T (100 mM Tris, pH 7.5, 0.9% NaCl, 0.05% Tween-20). Proteins were detected with primary antibodies listed after overnight incubation at 4°C. After washing 3x 5min with TBS-T, secondary anti-rabbit (Cell Signaling, #7074S) or anti-mouse HRP-linked antibodies (Cell Signaling, #7076S) were incubated for 1h at RT. Images were acquired with WesternBright Sirius Chemiluminescent Detection Kit (Advansta) in Fusion FX7 imaging system (Vilber).

### Co-Immunoprecipitation

For whole cell lysates, cells were lysed with RIPA without SDS with Halt protease inhibitor and phosphatase inhibitor cocktail (Thermo Scientific, #78420) for 30min on ice, as described above. For the isolation of the nuclear fraction, cells were collected and crosslinked with 0.1% PFA-PBS for 8min at RT. The reaction was quenched with 125 mM Glycine, cells were washed with PBS. The cell pellet was resuspended in ice-cold fractionation buffer (0.5 M sucrose, 50 mM KCl, 5 mM MgCl2, 50 mM Tris–HCl, pH 7.4) and incubated on ice. After 5min, NP-40 was added to a final concentration of 0.1% and samples were vortexed. Nuclei were collected by centrifugation at 3300 rcf for 3min at 4°C. The pellet containing the nuclei was lysed on ice with RIPA buffer for 30min, samples were vortexed every 5min. Protein concentration was assessed with Pierce BCA protein Assay Kit (Thermo Scientific, #23225). For the pull-down, 50 µl of Dynabeads Protein G (Invitrogen, #10003D) were coated with 5 µg of TAOK1, TRIM25 or PCNA antibody for each sample. 1 to 1.5 mg of protein lysates were loaded to the beads and incubated rotating overnight at 4°C. For immunoblot analysis beads were washed 3 times with 0.02% PBS-T and eluted by boiling in 6x sample buffer at 95°C for 10 min. Protein was analyzed on SDS/PAGE as described above. For mass spectrometry analysis, beads were washed 5 times with TBS before on-bead digestion for mass spectrometry analysis.

### Mass spectrometry analysis of co-IP

Proteins trapped on IP beads were treated essentially as described elsewhere with some changes (*66*). The digests were acidified with trifluoroacetic acid and analyzed by nano– liquid chromatography (LC)–MS/MS (Ultimate 3000 coupled to a QExactive HF mass spectrometer, Thermo Fisher Scientific) with three injections of 5-μl digest each. Peptides were trapped on a micro-precolumn C18 PepMap100 (5 μm, 100 Å, 300 μm × 5 mm, Thermo Fisher Scientific, Reinach, Switzerland) and separated by backflush onto the analytical nano-column (C18, 3 μm, 155 Å, 0.075 mm inside diameter × 150 mm length, Nikkyo Technos, Tokyo, Japan) applying a 40-min gradient of 5% acetonitrile to 40% in water and 0.1% formic acid at a flow rate of 350 nl/min. With two injections the data was acquired in data-dependent mode as described earlier. The third injection was used for a data-independent acquisition strategy isolating precursors in the range of m/z 330-1600 with 14 isolation windows of variable size. The full-scan method was set with resolution at 120,000 with an automatic gain control (AGC) target of 3 × 10^6^ and maximum ion injection time of 60 ms. The data-independent method for precursor ion fragmentation was applied with the following settings: resolution 30,000, AGC of 3 × 10^6^, maximum ion time of 47 ms, stepped collision energy of 25.5-27.0-30.0, respectively.

MS data were interpreted with fragpipe using the built-in ion quant algorithm for MS1-based ion quantification (version 22.0) (*67*) against a forward and reversed Swiss-Prot mouse protein sequence fasta file (release 2024_02) using the following settings: strict trypsin cleavage rule allowing up to three missed cleavage sites; variable modifications of acetylated protein N termini, oxidation of methionine, and pyro-Glu from Glu and Asp, as well as fixed carbamidomethylation of cysteines; peptide and fragment mass tolerances of 10 and 20 parts per million (ppm); no match between runs, respectively. Identified peptides and proteins were filtered to a 1% false discovery rate based on reversed database matches, and a minimum of two peptides were requested for acceptance of a protein group identification.

### Immunofluorescence

Cells were grown on coverslips in 24-well plates 2 days prior to experiment. To analyze RAD51 and RPA foci, cells were treated with 10 µM olaparib overnight or irradiated with 10 Gy and fixed 3 h post IR. For micronuclei staining cells were treated with 0.5 µM olaparib for 48 h prior fixation. Fixation was performed with 4% paraformaldehyde for 20 min on ice. Fixed cells were washed with PBS and permeabilized with 0.2% (v/v) Triton X-100 for 20min. After permeabilization, slides were blocked with staining buffer (PBS, BSA (2% w/v), glycine (0.15% w/v), Triton X-100 (0.1% v/v)) for 1h at RT. Primary antibodies were incubated at the indicated dilution in staining buffer for 1 h at RT or at 4°C overnight if used for PLA/SIRF assay. Slides for simple immunofluorescence were then washed 3 times with staining buffer and incubated with goat anti-rabbit IgG (H+L) Cross-Absorbed Secondary Antibody, Texas Red (Thermo Scientific, #T-2767) or Alexa-Fluor 488 (Thermo Scientific, #A-11008) or goat anti-mouse IgG (H+L) Cross-Absorbed Secondary Antibody, Texas Red (Thermo Scientific, #T-862) or Alexa-Fluor 488 (Thermo Scientific, #A-11029) diluted 1:2500 in staining buffer and DAPI (Thermo Scientific, #D21490, 1:50,000) for 1 h at RT. Slides were washed 5 times with staining buffer before mounting with Dako Fluorescence Mounting Medium (Agilent Technologies, #S3023). Images were acquired with Delta Vision Elite widefield microscope (GE Healthcare Life Sciences) or Nikon Ti2 Cicero spinning disc (Nikon). Multiple fields of view were imaged per sample with 60x objective at the resolution of 2048 x 2048 pixels. Image analysis was performed using Fiji image processing package of ImageJ or NIS Elements General Analysis 3 tool (Nikon).

For SIRF assay cells were labelled with EdU for 10 min (25 µM), washed and if indicated incubated with 2 mM HU for 2 h. Slides for PLA/SIRF assay were pre-extracted for 5 min on ice in 10 mM PIPES pH 7, 0.3 M sucrose, 0.1 M NaCl, 3 mM MgCl_2_ before fixation. Fixation and permeabilization was performed as described above. SIRF samples were processed with click reaction (100 mM Tris pH 8, 100 mM CuSO4, 2 mg/ml sodium-L-ascorbate, 10 mM biotin-azide) for 120 min at 37 °C. Slides were then blocked and incubated with primary antibody as indicated above. After primary antibody, cells were washed 2 times 5 min with PLA buffer A (Duolink kit, Sigma, #DUO92101) and incubated 1 h at 37°C with PLA probes. Next slides were washed again 2 times 5 min with buffer A before incubation with ligase for 30min at 37°C followed by amplification for 100 min at 37°C. Before incubating with secondary antibody and DAPI (as described above), slides were washed with PLA buffer B twice for 10 min. Slides were mounted and acquired as described above.

### DNA fiber analysis

Fork progression was measured as previously described (*68*). In brief, asynchronously growing KB1P-G3B1, KB1P-G3 or KB2P-1.21 cells were labeled with 25 µM thymidine analogue 5-chloro-2’-deoxyuridine (CIdU) (Sigma-Aldrich, #C6891) for 20 min, washed 3 times with PBS followed by labeling with 250 µM thymidine analogue 5-iodo-2′-deoxyuridine (IdU) for 20 min. Cells were either untreated or treated with 4 mM HU for 3 h. Cells were collected by trypsinization and 2 µl of the cell suspension was lysed with 8 µl lysis buffer (200 mM Tris-HCl, pH 7.4, 50 mM EDTA, and 0.5% (v/v) SDS) on a positively charged microscope slide. After 8 min of incubation, slides were tilted at an ∼30° angle to spread DNA fibers. Slides were then fixed with methanol: acetic acid (3:1) for 10 min at RT. After 2 times 3 min wash with PBS, slides were incubated with HCl (2.5 M) for 1 h at RT, washed 5 times with PBS and blocked for 40 min in 2% (w/v) BSA in 0.1% (v/v) PBS-T (PBS and Tween 20). CldU and IdU tracks were stained with primary antibodies anti-BrdU/CldU and anti-BrdU/IdU for 3 h at RT. After 5 times 3 min wash with PBS-T slides were stained with goat anti-mouse IgG (H+L) Cross-Absorbed Secondary Antibody, Alexa-Fluor 488 (Thermo Scientific, #A-11029, 1:300) and Alexa-Fluor 594 goat anti-rat IgG (Thermo Scientific, #A11007, 1:300) for 1 h at RT. Finally, slides were washed 5 times 3 min with PBS-T and mounted with Fluorescence mounting media (Dako). Images were acquired with Leica DMI4000B microscope at 60x magnification. Fiber tracks were measured manually using Fiji software and segmented line tool.

ssDNA gap formation was measured as previously described (*68*). In brief, asynchronously growing KB1P-G3 cells were exposed to 10 µM olaparib 2 h prior labelling. Cells were labelled for 10 min with 25 µM CldU followed by 45 min 250 µM IdU in media with or without 10 µM olaparib. After collection, cells were pre-extracted (100 mM NaCl, 10 mM MOPS pH 7, 3 mM MgCl2, 300 mM sucrose and 0.5% Triton X-100 in water) for 10 min at RT. Nuclei were collected by centrifugation at 8000 rpm for 5 min. Next, nuclei were incubated for 30 min at 37°C in S1 nuclease buffer with or without 20 U/ml S1 nuclease (Thermo Scientific, #EN0321). The process was quenched after 30 min with PBS + 0.1% BSA and fiber spreading and staining was performed as described above.

### Native BrdU assay

Experiment was carried out as described previously (*69*) with modifications. Cells were labelled with 10 µM CldU for 48 hours. On the day of the experiment, cells were treated with PARP inhibitors (KB1P-G3: 0.75 µM for 5 h, and MDA-MB-436: 1 µM for 8 h) and pre-extracted with ice-cold CSK buffer (10 mM PIPES, 100 mM NaCl, 300 mM Sucrose, 250 mM EGTA, 1 mM MgCl2, 1 mM DTT and protease inhibitor cocktail) on ice for 5 minutes. Cells were then fixed with pre-warm 4% formaldehyde for 15 minutes in 37°C and permeabilized with 0.1% Triton X-100 in RT. Permeabilized cells were blocked with 5% BSA in PBS and incubated with anti-BrdU antibody (Abcam ab6326) at 1:200 dilution for 1 h in RT. Cells were then washed and incubated with anti-rat Alexa FluorTM 488 at 1:1000 dilution for 1 h in RT. After washing, cells were stained with 0.1 µg/ml DAPI for 15 minutes. High content automated imaging was performed using Opera Phenix confocal spinning disk microscope with 3 Z stacks per field (1 µm/stack). Maximum intensity projection images were used for quantifying the intensities of each channel using CellProfiler software 4.2.1.

### DNA fibers analysis with S1 nuclease

Cells were seeded on coverslips for 24 h and labelled sequentially with 25 µM CldU for 15 minutes and 250 µM IdU with or without 10 µM PARP inhibitors for 45 minutes. Immediately after the treatment, cells were pre-extracted (100 mM NaCl, 10 mM MOPS pH 7, 3 mM MgCl2, 300 mM sucrose and 0.5% Triton X-100 in water) for 10 minutes in RT. Pre-extracted cells were gently washed with PBS and incubated with 20 U/ml S1 nuclease (Thermo Scientific, #EN0321) for 30 minutes at 37°C. After S1 nuclease treatment coverslips were processed for lysing and spreading using the combing machine designed and manufactured by the Department of Experimental Medical Instrumentation (EMI), Erasmus MC with the retraction rate of 300 μm per second (*70*). Coverslips were air-dried and fixed with methanol: acetic acid (3:1) at 4°C overnight, denatured with 2.5 M HCl at RT for 1 hour, and immunolabelled as described previously (*69*).

### DSB-Spectrum reporter

DSB-Spectrum_V1 reporter assay was performed as previously described (*40*) with few adaptations. HEK293FT cells containing DSB-Spectrum_V1 were a kind gift from Michael B. Yaffe and Haico van Attikum. Briefly, 10,000 cells were seeded per well in a 96-well plate on day 0, cells were seeded in quadruplicates for each siRNA. On day 1, transfection with the indicated siRNA was performed with Lipofectamine RNAiMAX (Thermo Scientific, #13778150) according to manufacturer’s protocol. On day 2, after 24 h media was refreshed. After 6h, duplicates of target cells were transfected with pX459-Cas9-sgRNA-iRFP constructs containing either an AAVS1 sgRNA (control) or an sgRNA targeting DSB-Spectrum BFP sequence. Transfection was performed with Lipofectamine 2000 (Thermo Scientific, #11668030). On day 6, cells were collected and subjected to flow cytometry with the CytoFLEX S (Beckman Coulter) for RFP, GFP and BFP intensity. Data was analyzed with FlowJo software. Fractions of HR positive (GFP+) or NHEJ positive (BFP+) population were calculated from cells transfected with the sgRNA construct (RFP+).

### *In vitro* kinase assay

Confluent monolayers of cells expressing HA-tagged wild type TAOK1 and mutants were harvested from T25 tissue culture flasks and cell pellets were lysed in 20 mM Tris pH 7.5, 140 mM KCl,1.8 mM MgCl_2_, 0.5% NP40, supplemented with EDTA-free protease inhibitor cocktail (Roche) on ice for 1 h with vortexing briefly every 10 minutes. Lysate was clarified by centrifugation at 16,000 g on a microfuge and soluble supernatant was used for immunoprecipitation with Pierce Anti-HA Magnetic Beads (Thermo Fisher Scientific, #88836).

On bead kinase assay was performed in a buffer containing 20 mM Tris pH 7.5, 10 mM MgCl_2,_ 0.1 mg/ml BSA, 40 µM myelin basic protein and 1 µM ATP, at 37°C for 1 h. Then reaction was separated from the beads and remaining ATP in the assay buffer was measured using Kinase-Glo® Luminescent Kinase Assays (Promega). Reaction mixture was transferred to 96 well plates allowed to equilibrate to room temperature for 20 min and luminescence counts were measured using Hidex plate reader.

### Immunohistochemistry

FFPE tissue blocks were collected from tissue archive at the Institute of Tissue Medicine and Pathology, University of Bern, Switzerland. Tumor tissue was selected from patinets from Inselspital Bern with signed general consent. Staining was performed with a BOND RX staining platform including a ready-to-use BOND Polymer Refine Detection Kit (DS9800). Tissue sections were dried (60°C, 30min), deparaffinized, PO blocked, detected and counterstained with Haematoxilin using DS9800 detection kit. Pre-treatment was done with Tris-EDTA buffer for 20 min at 95°C and stained with TAOK1 antibody (Proteintech, #26250-1-AP, 1:100) for 30 min at RT.

### SCAN-B Survival Analysis

Survival analysis for TAOK1 high/low expression was performed on the publicly available SCAN-B database, accessible under GEO accession number GSE96058. Gene expression and survival data were incorporated and Kaplan-Meier survival curves were plotted using R software (version 4.4.3) and package ggsurvfit. Either all patients in the database were included in the survival analysis or patients were selected according to chemotherapy/non-chemotherapy treated, and two groups (high / low expression) were established using the median expression as a cutoff. Log-rank test was then used to determine statistical significance.

### Statistical Analysis

Statistical analysis was performed using GraphPad Prism 10 or R software. Significant differences are indicated with p-values. P-values above 0.05 were considered non-significant and are indicated with “ns”. Applied statistical tests of the figures are indicated in the corresponding figure legends.

## Acknowledgments

We would like to thank Hana Hanzlikova, Diego Dibitetto and Carmen Perry for critical reading of the manuscript and all members of the Rottenberg lab for helpful discussions. We thank the Translational Research Unit TRU of the University of Bern for helping with IHC sample collection and stainings.

## Funding

Swiss National Science Foundation grant 320030M_219453 (JJ, SR)

European Research Council grant ERC-2019-AdG-883877 (SR)

Swiss Cancer Research Foundation grant KFS-5519-02-2022 (SR)

Office of the Assistant Secretary of Defense for Health Affairs through the Ovarian Cancer Research Program under Award No. W81XWH-22-1-0557 (SR)

ISREC Foundation grant (SR)

Foundation for Clinical-Experimental Cancer Research grant (PF)

## Author contributions

Conceptualization: LL, PF, SR

Funding acquisition: PF, SR

Methodology: LL, PF, SR

Investigation: LL, MM, AN, IK, EG, DAS, PMS, SH, RK, CF, ME, MGF, EV, SdB

Visualization: LL, MGF

Supervision: PF, SR, MvV, ARC, JJ

Writing—original draft: LL, PF, SR

## Competing interests

All other authors declare they have no competing interests.

## Data and materials availability

All data supporting the findings of this study are included in the main text or the supplementary materials. Additional data may be requested from the authors.

## Supplementary Materials

**fig. S1.**
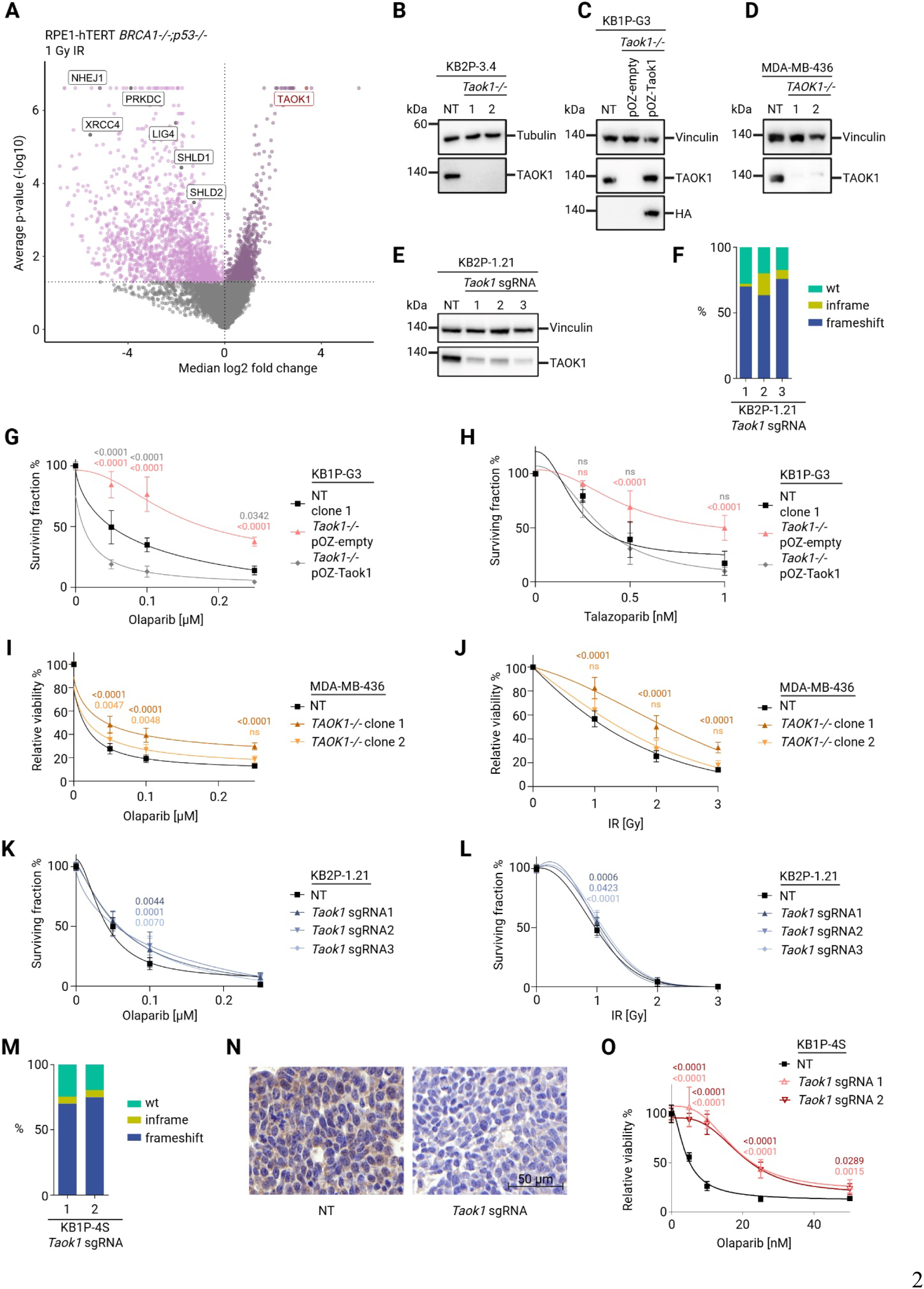
**(A)** Volcano plot representing gene summary of MAGeCK analysis of the whole genome CRISPR-Cas9 screens in RPE1-hTERT *BRCA1^-/-^;p53^-/-^* cells treated with 1 Gy IR. (**B-E**) Western blot analysis of TAOK1 protein expression of NT and TAOK1 KO in KB2P-3.4 (B), KB1P-G3 including HA-tagged, *Taok1* cDNA rescue (pOZ-Taok1) (C), MDA-MB-436 (D) and KB2P-1.21 (E) cell lines. (**F**) TIDE analysis of polyclonal KB2P-1.21 cells. (**G, H**) Growth assays in KB1P-G3 NT, *Taok1^-/-^* and *Taok1* rescue cells treated with the indicated doses of olaparib (G) or talazoparib (H). Statistical analysis was performed using two-way ANOVA followed by Dunnett’s test. (**I, J**) Growth assays in MDA-MB-436 NT and *TAOK1*^-/-^ cell lines treated with indicated doses of olaparib (I) or IR (J). Statistical analysis was performed using two-way ANOVA followed by Dunnett’s test. (**K, L**) Growth assays in KB2P-1.21 NT and *Taok1* sgRNA cells treated with the indicated doses of olaparib (K) or IR (L). Statistical analysis was performed using two-way ANOVA followed by Dunnett’s test. (**M**) TIDE analysis of polyclonal KB1P-4S organoids. (**N**) Immunohistochemistry staining for TAOK1 of untreated BRCA1;p53-deficient mouse mammary tumors with NT or *Taok1* sgRNA. (**O**) Olaparib response of KB1P-4S organoids transduced with NT or *Taok1* sgRNAs. Statistical analysis was performed using two-way ANOVA followed by Dunnett’s test.

**fig. S2.**
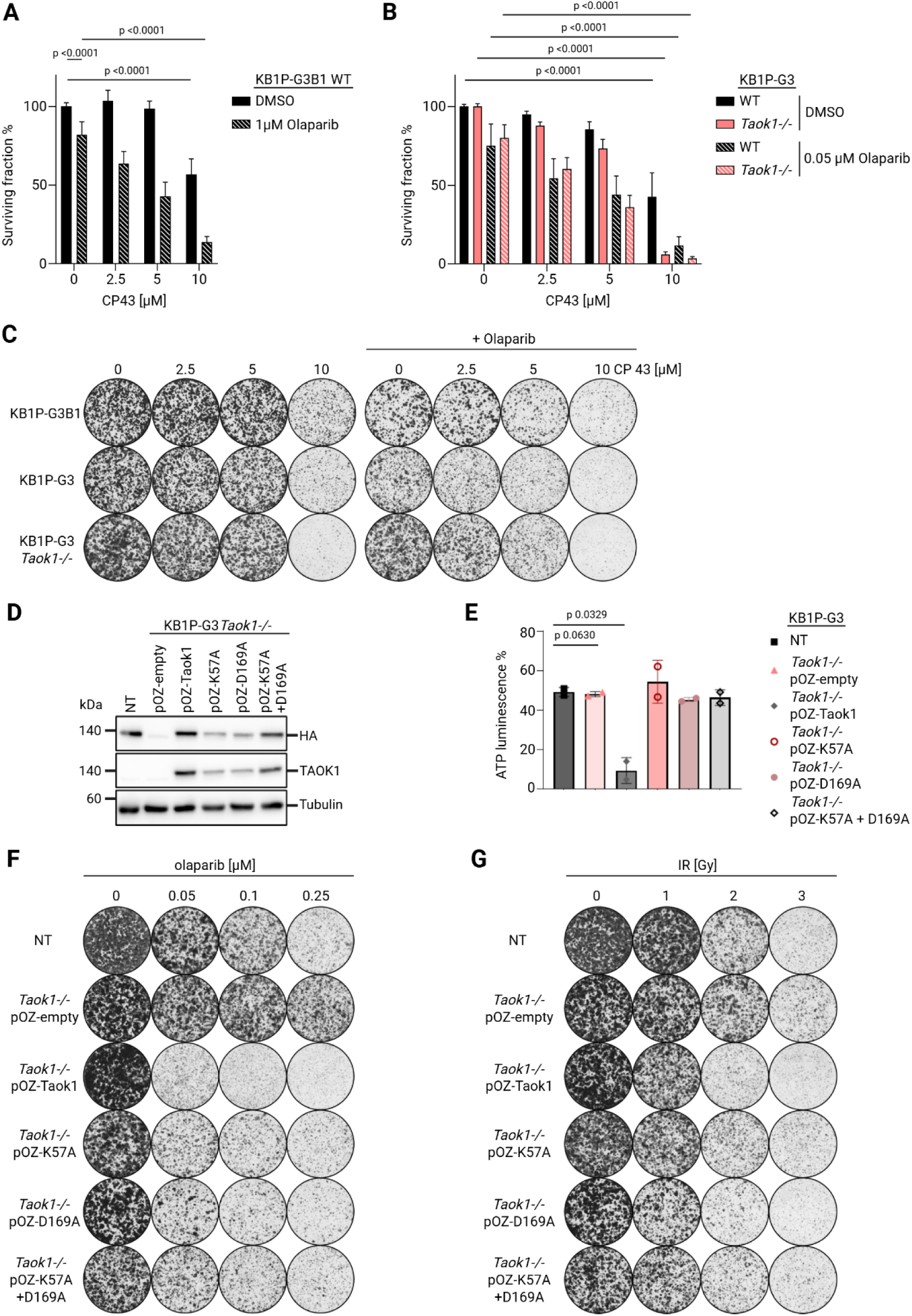
(**A-C**) Quantification (A, B) and representative images (C) of growth assays in KB1P-G3B1 WT (A) and KB1P-G3 WT and *Taok1^-/-^* cells treated with the indicated doses of CP 43 and olaparib. Statistical analysis was performed using two-way ANOVA followed by Dunnett’s test. (**D**) Western blot analysis of expression level of HA-tagged *Taok1* rescue constructs. (**E**) *In vitro* kinase assay of indicated KB1P-G3 cell lines. Protein was isolated from KB1P-G3 cell lines by co-IP with HA-tag. NT cell line was used as negative control with no HA-tag while the other cell lines expressed an HA-tag with the indicated construct. Bars represent mean ± SD, statistical analysis was done with two-way ANOVA followed by Dunnett’s test. (**F, G**) Representative images of growth assay of KB1P-G3 NT, *Taok1^-/-^*and *Taok1* full cDNA rescue (pOZ-Taok1) or two different kinase mutants (pOZ-K57A, pOZ-D169A) and a double mutant (pOZ-K57A+D169A) treated with indicated doses of olaparib (F) or IR (G).

**fig. S3.**
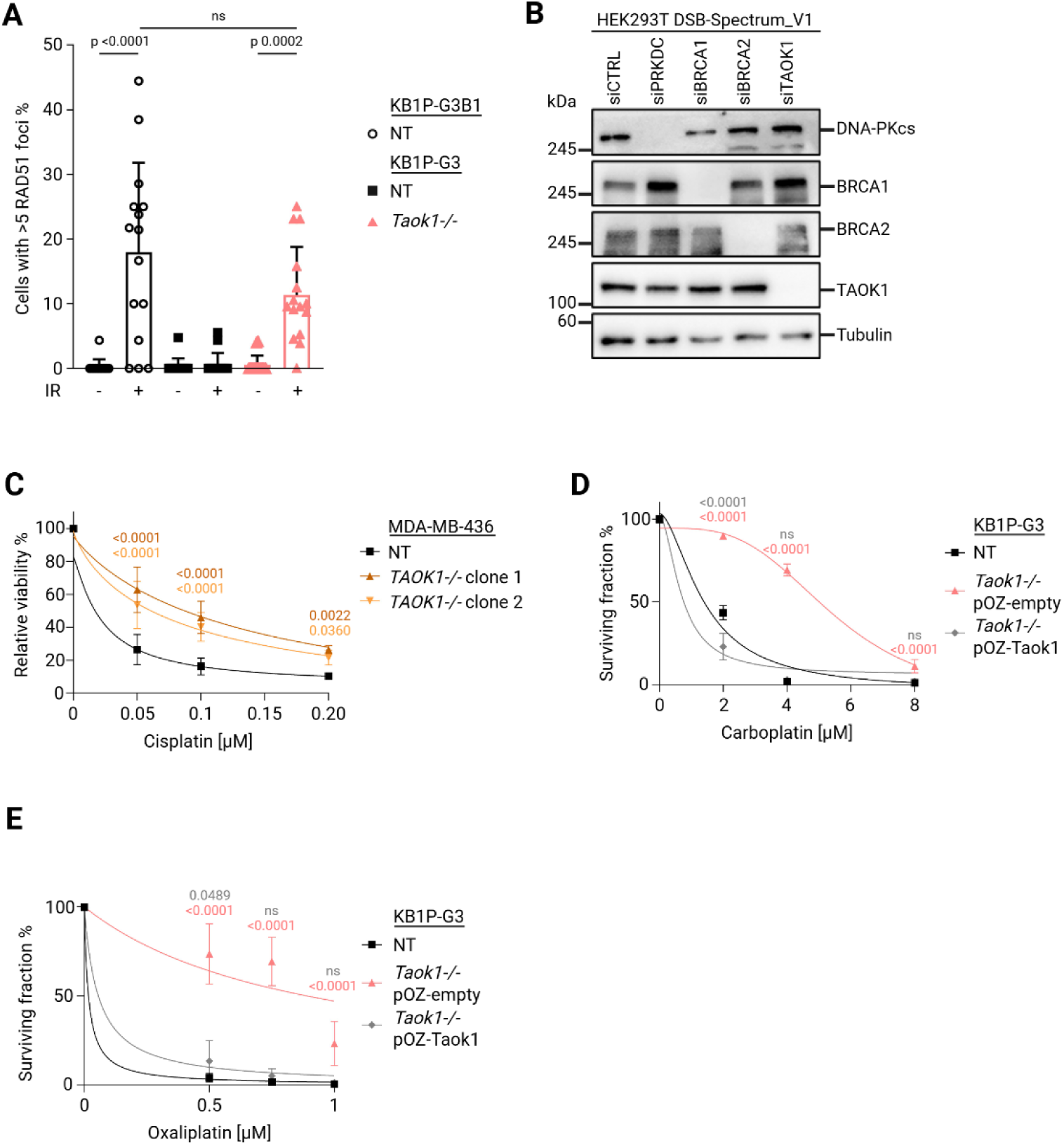
(**A**) Quantification of RAD51 foci formation 3 h post 10 Gy IR. Bars represent mean ± SD, statistical analysis was performed using ANOVA followed by Tukey’s multiple comparison test. (**B**) Verification of siRNA KD efficiency in HEK293T DSB-Spectrum_V1 cell line. (**C**) Growth assays with MDA-MB-436 NT and TAOK1 KO cell lines treated with the indicated doses of cisplatin. Statistical analysis was performed using two-way ANOVA followed by Dunnett’s test. (**D, E**) Growth assays in KB1P-G3 NT, TAOK1 KO and *Taok1* rescue cell lines treated with indicated doses of carboplatin (D) or oxaliplatin (E). Statistical analysis was performed using two-way ANOVA followed by Dunnett’s test.

**fig. S4.**
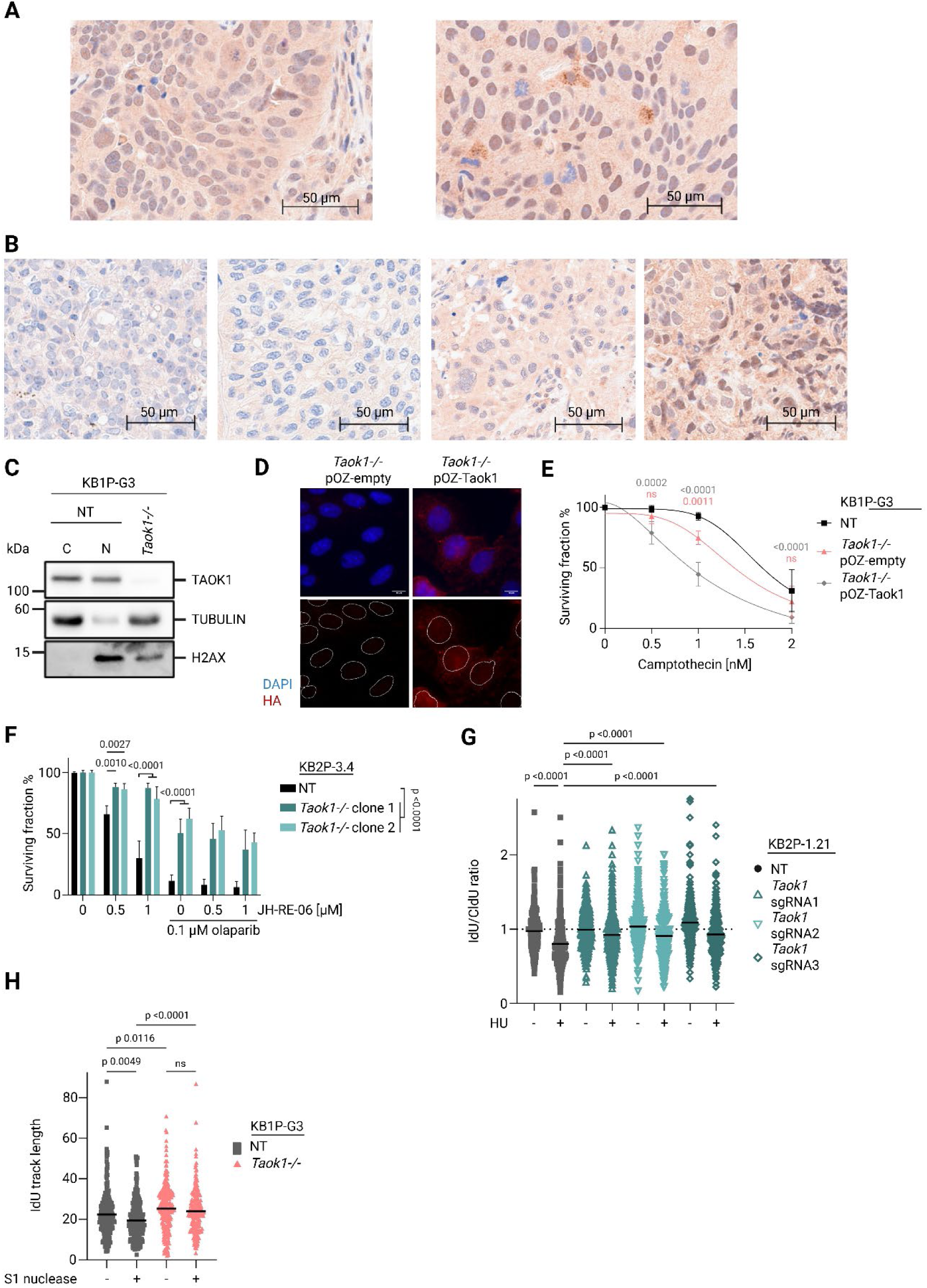
**(A)** Representative images of IHC staining for TAOK1 on human tumor samples of prostate and lung showing variable nuclear and diffuse cytoplasmatic localization. (**B**) Representative images of IHC staining for TAOK1 on human tumor samples of prostate, bladder and ovary showing varying TAOK1 expression levels. (**C**)Western blot analysis of KB1P-G3 NT samples of cytoplasmatic (C) and nuclear (N) fraction and *Taok1^-/-^* whole cell lysate. (**D**) Representative images of immunofluorescence for cellular expression pattern of KB1P-G3 *Taok1^-/-^* cells with HA-tagged pOZ-Taok1 rescue construct compared to pOZ-empty vector control. (**E**) Growth assay in KB1P-G3 NT, *Taok1^-/-^* and *Taok1* rescue cell lines treated with indicated doses of camptothecin. Statistical analysis was performed using two-way ANOVA followed by Dunnett’s test. (**F**) Growth assay in KB2P-3.4 NT and *Taok1^-/-^*clones treated with indicated doses of JH-RE-06 with or without olaparib (0.1 µM). Statistical analysis was performed using two-way ANOVA followed by Tukey’s test. (**G**) DNA fiber analysis in indicated cell lines with or without 4 mM HU treatment (3 h). Bar represents mean of IdU/CldU ratio of at least 250 fibers, statistical analysis was done with ANOVA followed by Tukey’s multiple comparison test. **(H)** DNA fiber analysis in indicated cell lines treated with 10 µM olaparib 2h prior and while labelling with CldU and IdU, ssDNA was digested with S1 nuclease if indicated. Bar represents the mean of IdU track length of at least 250 fibers, statistical analysis was done with ANOVA followed by Tukey’s multiple comparison test.

**fig. S5.**
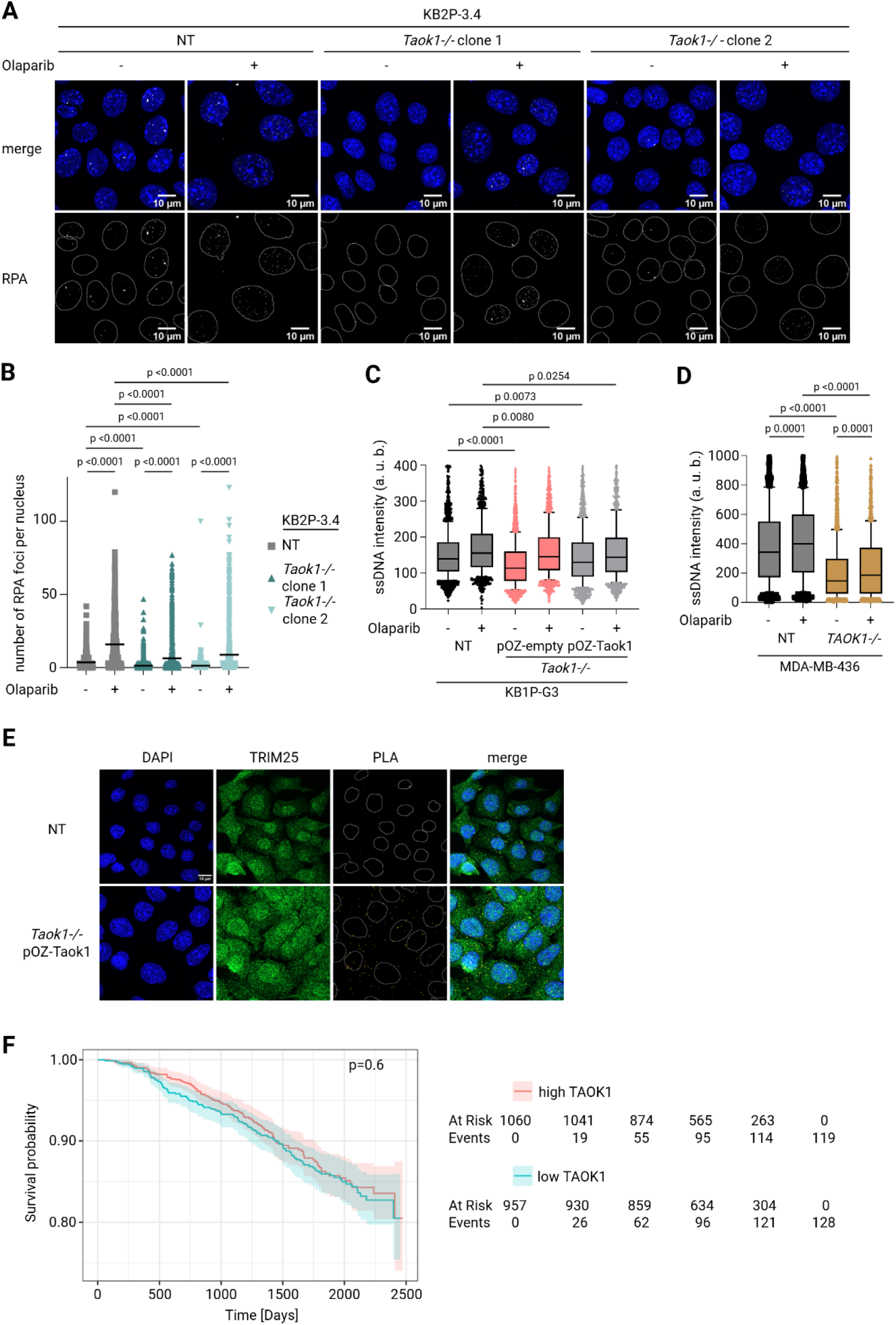
(**A, B**) Representative images (A) and quantification (B) of immunofluorescence analysis of RPA foci formation in KB2P-3.4 NT and *Taok1^-/-^*cells upon olaparib treatment (10 µM for 16 h). Bars indicate mean number of foci per nucleus of at least 1000 nuclei. Statistical analysis was done with ANOVA followed by Tukey’s multiple comparison test. (**C, D**) Graph representing ssDNA intensity of native BrdU incorporation in KB1P-G3 (C) and MDA-MB-436 (D) cell lines. Cells were treated with 0.75 µM olaparib for 5 h. Ordinary one-way ANOVA with Dunnett’s multiple comparison test was used for statistical analysis. (**E**) Representative images of PLA assay between TRIM25 and HA-tag in KB1P-G3 NT and *Taok1* rescue cells. (**F**) Kaplan-Meier curve of survival probability of SCAN-B dataset treated with non-chemotherapy. Statistical analysis was done with log-rank test.

**Data S1. (separate file)**

Table of gene summary of MAGeCK analysis of the kinome shRNA screens in KB2P-1.21 cells treated with olaparib. Log2FC values were calculated between untreated control and olaparib treated samples. Results were filtered for >3 and <10 good_sgRNA.

**Data S2. (separate file)**

Table of gene summary of MAGeCK analysis of the Brunello whole genome CRISPR-Cas9 screens in RPE1-hTERT *BRCA1^-/-^;p53^-/-^* cells treated with indicated dose of IR. Log2FC values were calculated between RT treated samples and Day 0. Results were filtered for >3 and <6 good_sgRNA.

**Data S3. (separate file)**

Table of mass spectrometry analysis of coIP for TAOK1, comparing pulldown of KB1P-G3 NT and TAOK1 KO. Log2FC values are the result of 4 independent replicas and were calculated with fragpipe.

## Notes

### Competing Interest Statement

The authors have declared no competing interest.

## References

1. R. Prakash, Y. Zhang, W. Feng, M. Jasin, Homologous Recombination and Human Health: The Roles of BRCA1, BRCA2, and Associated Proteins. Cold Spring Harb. Perspect. Biol. 7, a016600 (2015).

2. J. A. Nickoloff, D. Jones, S.-H. Lee, E. A. Williamson, R. Hromas, Drugging the Cancers Addicted to DNA Repair. JNCI J. Natl. Cancer Inst. 109 (2017).

3. C. Kan, J. Zhang, BRCA1 Mutation: A Predictive Marker for Radiation Therapy? *Int*. J. Radiat. Oncol. 93, 281–293 (2015).

4. H. Gaillard, T. García-Muse, A. Aguilera, Replication stress and cancer. Nat. Rev. Cancer 15, 276–289 (2015).

5. T. Ubhi, G. W. Brown, Exploiting DNA Replication Stress for Cancer Treatment. Cancer Res. 79, 1730–1739 (2019).

6. H. Zhu, U. Swami, R. Preet, J. Zhang, Harnessing DNA Replication Stress for Novel Cancer Therapy. Genes 11, 990 (2020).

7. M. Petropoulos, A. Karamichali, G. G. Rossetti, A. Freudenmann, L. G. Iacovino, V. S. Dionellis, S. K. Sotiriou, T. D. Halazonetis, Transcription–replication conflicts underlie sensitivity to PARP inhibitors. Nature 628, 433–441 (2024).

8. J. Murai, S. N. Huang, B. B. Das, A. Renaud, Y. Zhang, J. H. Doroshow, J. Ji, S. Takeda, Y. Pommier, Trapping of PARP1 and PARP2 by Clinical PARP Inhibitors. Cancer Res. 72, 5588–5599 (2012).

9. K. Schlacher, N. Christ, N. Siaud, A. Egashira, H. Wu, M. Jasin, Double-Strand Break Repair-Independent Role for BRCA2 in Blocking Stalled Replication Fork Degradation by MRE11. Cell 145, 529–542 (2011).

10. K. Schlacher, H. Wu, M. Jasin, A Distinct Replication Fork Protection Pathway Connects Fanconi Anemia Tumor Suppressors to RAD51-BRCA1/2. Cancer Cell 22, 106–116 (2012).

11. A. Ray Chaudhuri, E. Callen, X. Ding, E. Gogola, A. A. Duarte, J.-E. Lee, N. Wong, V. Lafarga, J. A. Calvo, N. J. Panzarino, S. John, A. Day, A. V. Crespo, B. Shen, L. M. Starnes, J. R. D. Ruiter, J. A. Daniel, P. A. Konstantinopoulos, D. Cortez, S. B. Cantor, O. Fernandez-Capetillo, K. Ge, J. Jonkers, S. Rottenberg, S. K. Sharan, A. Nussenzweig, Replication fork stability confers chemoresistance in BRCA-deficient cells. Nature 535, 382–387 (2016).

12. P. X. Lim, M. Zaman, W. Feng, M. Jasin, BRCA2 promotes genomic integrity and therapy resistance primarily through its role in homology-directed repair. Mol. Cell 84, 447–462.e10 (2024).

13. D. Dibitetto, C. A. Widmer, S. Rottenberg, PARPi, BRCA, and gaps: controversies and future research. Trends Cancer 10, 857–869 (2024).

14. D. Dibitetto, M. Liptay, F. Vivalda, H. Dogan, E. Gogola, M. González Fernández, A. Duarte, J. A. Schmid, M. Decollogny, P. Francica, S. Przetocka, S. T. Durant, J. V. Forment, I. Klebic, M. Siffert, R. De Bruijn, A. N. Kousholt, N. A. Marti, M. Dettwiler, C. S. Sørensen, J.-C. Tille, M. Undurraga, I. Labidi-Galy, M. Lopes, A. A. Sartori, J. Jonkers, S. Rottenberg, H2AX promotes replication fork degradation and chemosensitivity in BRCA-deficient tumours. Nat. Commun. 15, 4430 (2024).

15. K. Pappas, M. Ferrari, P. Smith, S. Nandakumar, Z. Khan, S. B. Young, J. LaClair, M. V. Russo, E. Huang-Hobbs, N. Schultz, W. Abida, W. Karthaus, M. Jasin, C. L Sawyers, BRCA2 reversion mutation-independent resistance to PARP inhibition through impaired DNA prereplication complex function. Proc. Natl. Acad. Sci. U. S. A. 122, e2426743122 (2025).

16. C. Hoege, B. Pfander, G.-L. Moldovan, G. Pyrowolakis, S. Jentsch, RAD6-dependent DNA repair is linked to modification of PCNA by ubiquitin and SUMO. Nature 419, 135–141 (2002).

17. M. Shaheen, I. Shanmugam, R. Hromas, The Role of PCNA Posttranslational Modifications in Translesion Synthesis. J. Nucleic Acids 2010, 761217 (2010).

18. J. Zámborszky, B. Szikriszt, J. Z. Gervai, O. Pipek, Á. Póti, M. Krzystanek, D. Ribli, J. M. Szalai-Gindl, I. Csabai, Z. Szallasi, C. Swanton, A. L. Richardson, D. Szüts, Loss of BRCA1 or BRCA2 markedly increases the rate of base substitution mutagenesis and has distinct effects on genomic deletions. Oncogene 36, 746–755 (2017).

19. T. Thakar, W. Leung, C. M. Nicolae, K. E. Clements, B. Shen, A.-K. Bielinsky, G.-L. Moldovan, Ubiquitinated-PCNA protects replication forks from DNA2-mediated degradation by regulating Okazaki fragment maturation and chromatin assembly. Nat. Commun. 11, 2147 (2020).

20. J. M. Park, S. W. Yang, K. R. Yu, S. H. Ka, S. W. Lee, J. H. Seol, Y. J. Jeon, C. H. Chung, Modification of PCNA by ISG15 Plays a Crucial Role in Termination of Error-Prone Translesion DNA Synthesis. Mol. Cell 54, 626–638 (2014).

21. C. P. Wardlaw, J. H. J. Petrini, ISG15 conjugation to proteins on nascent DNA mitigates DNA replication stress. Nat. Commun. 13, 5971 (2022).

22. J. E. Jaspers, A. Kersbergen, U. Boon, W. Sol, L. van Deemter, S. A. Zander, R. Drost, E. Wientjens, J. Ji, A. Aly, J. H. Doroshow, A. Cranston, N. M. B. Martin, A. Lau, M. J. O’Connor, S. Ganesan, P. Borst, J. Jonkers, S. Rottenberg, Loss of 53BP1 Causes PARP Inhibitor Resistance in *Brca1* -Mutated Mouse Mammary Tumors. Cancer Discov. 3, 68–81 (2013).

23. J. Bhin, M. Paes Dias, E. Gogola, F. Rolfs, S. R. Piersma, R. de Bruijn, J. R. de Ruiter, B. van den Broek, A. A. Duarte, W. Sol, I. van der Heijden, C. Andronikou, T. S. Kaiponen, L. Bakker, C. Lieftink, B. Morris, R. L. Beijersbergen, M. van de Ven, C. R. Jimenez, L. F. A. Wessels, S. Rottenberg, J. Jonkers, Multi-omics analysis reveals distinct non-reversion mechanisms of PARPi resistance in BRCA1-versus BRCA2-deficient mammary tumors. Cell Rep. 42, 112538 (2023).

24. G. Xu, J. R. Chapman, I. Brandsma, J. Yuan, M. Mistrik, P. Bouwman, J. Bartkova, E. Gogola, D. Warmerdam, M. Barazas, J. E. Jaspers, K. Watanabe, M. Pieterse, A. Kersbergen, W. Sol, P. H. N. Celie, P. C. Schouten, B. van den Broek, A. Salman, M. Nieuwland, I. de Rink, J. de Ronde, K. Jalink, S. J. Boulton, J. Chen, D. C. van Gent, J. Bartek, J. Jonkers, P. Borst, S. Rottenberg, REV7 counteracts DNA double-strand break resection and affects PARP inhibition. Nature 521, 541–544 (2015).

25. S. M. Noordermeer, S. Adam, D. Setiaputra, M. Barazas, S. J. Pettitt, A. K. Ling, M. Olivieri, A. Álvarez-Quilón, N. Moatti, M. Zimmermann, S. Annunziato, D. B. Krastev, F. Song, I. Brandsma, J. Frankum, R. Brough, A. Sherker, S. Landry, R. K. Szilard, M. M. Munro, A. McEwan, T. Goullet de Rugy, Z.-Y. Lin, T. Hart, J. Moffat, A.-C. Gingras, A. Martin, H. van Attikum, J. Jonkers, C. J. Lord, S. Rottenberg, D. Durocher, The shieldin complex mediates 53BP1-dependent DNA repair. Nature 560, 117–121 (2018).

26. M. Barazas, S. Annunziato, S. J. Pettitt, I. de Krijger, H. Ghezraoui, S. J. Roobol, C. Lutz, J. Frankum, F. F. Song, R. Brough, B. Evers, E. Gogola, J. Bhin, M. van de Ven, D. C. van Gent, J. J. L. Jacobs, R. Chapman, C. J. Lord, J. Jonkers, S. Rottenberg, The CST Complex Mediates End Protection at Double-Strand Breaks and Promotes PARP Inhibitor Sensitivity in BRCA1-Deficient Cells. Cell Rep. 23, 2107–2118 (2018).

27. M. Barazas, A. Gasparini, Y. Huang, A. Küçükosmanoğlu, S. Annunziato, P. Bouwman, W. Sol, A. Kersbergen, N. Proost, R. de Korte-Grimmerink, M. van de Ven, J. Jonkers, G. R. Borst, S. Rottenberg, Radiosensitivity Is an Acquired Vulnerability of PARPi-Resistant BRCA1-Deficient Tumors. Cancer Res. 79, 452–460 (2019).

28. E. Gogola, A. A. Duarte, J. R. de Ruiter, W. W. Wiegant, J. A. Schmid, R. de Bruijn, D. I. James, S. Guerrero Llobet, D. J. Vis, S. Annunziato, B. van den Broek, M. Barazas, A. Kersbergen, M. van de Ven, M. Tarsounas, D. J. Ogilvie, M. van Vugt, L. F. A. Wessels, J. Bartkova, I. Gromova, M. Andújar-Sánchez, J. Bartek, M. Lopes, H. van Attikum, P. Borst, J. Jonkers, S. Rottenberg, Selective Loss of PARG Restores PARylation and Counteracts PARP Inhibitor-Mediated Synthetic Lethality. Cancer Cell 33, 1078–1093.e12 (2018).

29. J. C. Boggiano, P. J. Vanderzalm, R. G. Fehon, Tao-1 Phosphorylates Hippo/MST Kinases to Regulate the Hippo-Salvador-Warts Tumor Suppressor Pathway. Dev. Cell 21, 888–895 (2011).

30. C. L. C. Poon, J. I. Lin, X. Zhang, K. F. Harvey, The Sterile 20-like Kinase Tao-1 Controls Tissue Growth by Regulating the Salvador-Warts-Hippo Pathway. Dev. Cell 21, 896–906 (2011).

31. T.-C. Zhu, Z.-P. He, S.-T. Li, L. Zheng, X.-Y. Zheng, X.-L. Lan, C.-H. Qu, R.-C. Nie, C. Gu, L.-N. Huang, X.-X. Cai, Z.-C. Xiang, D. Xie, M.-Y. Cai, TAOK1 promotes filament formation in HR repair through phosphorylating USP7. Proc. Natl. Acad. Sci. 122, e2422262122 (2025).

32. K. R. Sanson, R. E. Hanna, M. Hegde, K. F. Donovan, C. Strand, M. E. Sullender, E. W. Vaimberg, A. Goodale, D. E. Root, F. Piccioni, J. G. Doench, Optimized libraries for CRISPR-Cas9 genetic screens with multiple modalities. Nat. Commun. 9, 5416 (2018).

33. W. Li, H. Xu, T. Xiao, L. Cong, M. I. Love, F. Zhang, R. A. Irizarry, J. S. Liu, M. Brown, X. S. Liu, MAGeCK enables robust identification of essential genes from genome-scale CRISPR/Cas9 knockout screens. Genome Biol. 15, 554 (2014).

34. H. Kim, E. George, R. L. Ragland, S. Rafail, R. Zhang, C. Krepler, M. A. Morgan, M. Herlyn, E. J. Brown, F. Simpkins, Targeting the ATR/CHK1 Axis with PARP Inhibition Results in Tumor Regression in *BRCA* - Mutant Ovarian Cancer Models. Clin. Cancer Res. 23, 3097–3108 (2017).

35. B. L. Mahaney, K. Meek, S. P. Lees-Miller, Repair of ionizing radiation-induced DNA double-strand breaks by non-homologous end-joining. Biochem. J. 417, 639–650 (2009).

36. F. Elstrodt, A. Hollestelle, J. H. A. Nagel, M. Gorin, M. Wasielewski, A. Van Den Ouweland, S. D. Merajver, S. P. Ethier, M. Schutte, *BRCA1* Mutation Analysis of 41 Human Breast Cancer Cell Lines Reveals Three New Deleterious Mutants. Cancer Res. 66, 41–45 (2006).

37. E. K. Brinkman, T. Chen, M. Amendola, B. van Steensel, Easy quantitative assessment of genome editing by sequence trace decomposition. Nucleic Acids Res. 42, e168–e168 (2014).

38. C. Zihni, C. Mitsopoulos, I. A. Tavares, A. J. Ridley, J. D. H. Morris, Prostate-derived Sterile 20-like Kinase 2 (PSK2) Regulates Apoptotic Morphology via C-Jun N-terminal Kinase and Rho Kinase-1. J. Biol. Chem. 281, 7317–7323 (2006).

39. M. Hutchison, K. S. Berman, M. H. Cobb, Isolation of TAO1, a Protein Kinase That Activates MEKs in Stress-activated Protein Kinase Cascades. J. Biol. Chem. 273, 28625–28632 (1998).

40. B. Van De Kooij, A. Kruswick, H. Van Attikum, M. B. Yaffe, Multi-pathway DNA-repair reporters reveal competition between end-joining, single-strand annealing and homologous recombination at Cas9-induced DNA double-strand breaks. Nat. Commun. 13, 5295 (2022).

41. R. Zellweger, D. Dalcher, K. Mutreja, M. Berti, J. A. Schmid, R. Herrador, A. Vindigni, M. Lopes, Rad51-mediated replication fork reversal is a global response to genotoxic treatments in human cells. J. Cell Biol. 208, 563–579 (2015).

42. S. Mijic, R. Zellweger, N. Chappidi, M. Berti, K. Jacobs, K. Mutreja, S. Ursich, A. Ray Chaudhuri, A. Nussenzweig, P. Janscak, M. Lopes, Replication fork reversal triggers fork degradation in BRCA2-defective cells. Nat. Commun. 8, 859 (2017).

43. S. Roy, J. W. Luzwick, K. Schlacher, SIRF: Quantitative in situ analysis of protein interactions at DNA replication forks. J. Cell Biol. 217, 1521–1536 (2018).

44. M. Raman, S. Earnest, K. Zhang, Y. Zhao, M. H. Cobb, TAO kinases mediate activation of p38 in response to DNA damage. EMBO J. 26, 2005–2014 (2007).

45. A. A. Davies, D. Huttner, Y. Daigaku, S. Chen, H. D. Ulrich, Activation of Ubiquitin-Dependent DNA Damage Bypass Is Mediated by Replication Protein A. Mol. Cell 29, 625–636 (2008).

46. M. R. Albertella, A. Lau, M. J. O’Connor, The overexpression of specialized DNA polymerases in cancer. DNA Repair 4, 583–593 (2005).

47. M. K. Zafar, R. L. Eoff, Translesion DNA Synthesis in Cancer: Molecular Mechanisms and Therapeutic Opportunities. Chem. Res. Toxicol. 30, 1942–1955 (2017).

48. J. Z. Gervai, J. Gálicza, Z. Szeltner, J. Zámborszky, D. Szüts, A genetic study based on PCNA-ubiquitin fusions reveals no requirement for PCNA polyubiquitylation in DNA damage tolerance. DNA Repair 54, 46–54 (2017).

49. D. Chen, J. Z. Gervai, Á. Póti, E. Németh, Z. Szeltner, B. Szikriszt, Z. Gyüre, J. Zámborszky, M. Ceccon, F. d’Adda Di Fagagna, Z. Szallasi, A. L. Richardson, D. Szüts, BRCA1 deficiency specific base substitution mutagenesis is dependent on translesion synthesis and regulated by 53BP1. Nat. Commun. 13, 226 (2022).

50. K. Cong, M. Peng, A. N. Kousholt, W. T. C. Lee, S. Lee, S. Nayak, J. Krais, P. S. VanderVere-Carozza, K. S. Pawelczak, J. Calvo, N. J. Panzarino, J. J. Turchi, N. Johnson, J. Jonkers, E. Rothenberg, S. B. Cantor, Replication gaps are a key determinant of PARP inhibitor synthetic lethality with BRCA deficiency. Mol. Cell 81, 3128–3144.e7 (2021).

51. R. N. Moro, U. Biswas, S. S. Kharat, F. D. Duzanic, P. Das, M. Stavrou, M. C. Raso, R. Freire, A. R. Chaudhuri, S. K. Sharan, L. Penengo, Interferon restores replication fork stability and cell viability in BRCA-defective cells via ISG15. Nat. Commun. 14, 6140 (2023).

52. W. Zou, D.-E. Zhang, The Interferon-inducible Ubiquitin-protein Isopeptide Ligase (E3) EFP Also Functions as an ISG15 E3 Ligase. J. Biol. Chem. 281, 3989–3994 (2006).

53. C. Brueffer, J. Vallon-Christersson, D. Grabau, A. Ehinger, J. Häkkinen, C. Hegardt, J. Malina, Y. Chen, P.-O. Bendahl, J. Manjer, M. Malmberg, C. Larsson, N. Loman, L. Rydén, Å. Borg, L. H. Saal, Clinical Value of RNA Sequencing-Based Classifiers for Prediction of the Five Conventional Breast Cancer Biomarkers: A Report From the Population-Based Multicenter Sweden Cancerome Analysis Network-Breast Initiative. *JCO Precis*. Oncol. 2, PO.17.00135 (2018).

54. L. H. Saal, J. Vallon-Christersson, J. Häkkinen, C. Hegardt, D. Grabau, C. Winter, C. Brueffer, M.-H. E. Tang, C. Reuterswärd, R. Schulz, A. Karlsson, A. Ehinger, J. Malina, J. Manjer, M. Malmberg, C. Larsson, L. Rydén, N. Loman, Å. Borg, The Sweden Cancerome Analysis Network - Breast (SCAN-B) Initiative: a large-scale multicenter infrastructure towards implementation of breast cancer genomic analyses in the clinical routine. Genome Med. 7, 20 (2015).

55. Z. Mirman, F. Lottersberger, H. Takai, T. Kibe, Y. Gong, K. Takai, A. Bianchi, M. Zimmermann, D. Durocher, T. De Lange, 53BP1–RIF1–shieldin counteracts DSB resection through CST- and Polα-dependent fill-in. Nature 560, 112–116 (2018).

56. H. Ghezraoui, C. Oliveira, J. R. Becker, K. Bilham, D. Moralli, C. Anzilotti, R. Fischer, M. Deobagkar-Lele, M. Sanchiz-Calvo, E. Fueyo-Marcos, S. Bonham, B. M. Kessler, S. Rottenberg, R. J. Cornall, C. M. Green, J. R. Chapman, 53BP1 cooperation with the REV7-shieldin complex underpins DNA structure-specific NHEJ. Nature 560, 122–127 (2018).

57. D. Vugic, I. Dumoulin, C. Martin, A. Minello, L. Alvaro-Aranda, J. Gomez-Escudero, R. Chaaban, R. Lebdy, C. Von Nicolai, V. Boucherit, C. Ribeyre, A. Constantinou, A. Carreira, Replication gap suppression depends on the double-strand DNA binding activity of BRCA2. Nat. Commun. 14, 446 (2023).

58. M. Dulovic-Mahlow, J. Trinh, K. K. Kandaswamy, G. J. Braathen, N. Di Donato, E. Rahikkala, S. Beblo, M. Werber, V. Krajka, Ø. L. Busk, H. Baumann, N. A. Al-Sannaa, F. Hinrichs, R. Affan, N. Navot, M. A. Al Balwi, G. Oprea, Ø. L. Holla, M. E. R. Weiss, R. A. Jamra, A.-K. Kahlert, S. Kishore, K. Tveten, M. Vos, A. Rolfs, K. Lohmann, De Novo Variants in TAOK1 Cause Neurodevelopmental Disorders. Am. J. Hum. Genet. 105, 213–220 (2019).

59. R. L. Shrestha, N. Tamura, A. Fries, N. Levin, J. Clark, V. M. Draviam, TAO1 kinase maintains chromosomal stability by facilitating proper congression of chromosomes. Open Biol. 4, 130108 (2014).

60. Y. Zheng, D. Pan, The Hippo Signaling Pathway in Development and Disease. Dev. Cell 50, 264–282 (2019).

61. T. Timm, X.-Y. Li, J. Biernat, J. Jiao, E. Mandelkow, J. Vandekerckhove, E.-M. Mandelkow, MARKK, a Ste20-like kinase, activates the polarity-inducing kinase MARK/PAR-1. EMBO J. 22, 5090–5101 (2003).

62. L. Chen, TAOK1 Promotes Proliferation and Invasion of Non-Small-Cell Lung Cancer Cells by Inhibition of WWC1. Comput. Math. Methods Med. 2022, 3157448 (2022).

63. B. Evers, R. Drost, E. Schut, M. De Bruin, E. Van Der Burg, P. W. B. Derksen, H. Holstege, X. Liu, E. Van Drunen, H. B. Beverloo, G. C. M. Smith, N. M. B. Martin, A. Lau, M. J. O’Connor, J. Jonkers, Selective Inhibition of BRCA2-Deficient Mammary Tumor Cell Growth by AZD2281 and Cisplatin. Clin. Cancer Res. 14, 3916–3925 (2008).

64. M. Olivieri, D. Durocher, Genome-scale chemogenomic CRISPR screens in human cells using the TKOv3 library. STAR Protoc. 2, 100321 (2021).

65. A. A. Duarte, E. Gogola, N. Sachs, M. Barazas, S. Annunziato, J. R De Ruiter, A. Velds, S. Blatter, J. M. Houthuijzen, M. Van De Ven, H. Clevers, P. Borst, J. Jonkers, S. Rottenberg, BRCA-deficient mouse mammary tumor organoids to study cancer-drug resistance. Nat. Methods 15, 134–140 (2018).

66. M. Stillinovic, M. A. Sarangdhar, N. Andina, A. Tardivel, F. Greub, G. Bombaci, C. Ansermet, M. Zatti, D. Saha, J. Xiong, T. Sagae, M. Yokogawa, M. Osawa, M. Heller, A. Keogh, I. Keller, A. Angelillo-Scherrer, R. Allam, Ribonuclease inhibitor and angiogenin system regulates cell type–specific global translation. Sci. Adv. 10, eadl0320 (2024).

67. F. Yu, S. E. Haynes, G. C. Teo, D. M. Avtonomov, D. A. Polasky, A. I. Nesvizhskii, Fast Quantitative Analysis of timsTOF PASEF Data with MSFragger and IonQuant. Mol. Cell. Proteomics 19, 1575–1585 (2020).

68. A. Quinet, D. Carvajal-Maldonado, D. Lemacon, A. Vindigni, “DNA Fiber Analysis: Mind the Gap!” in Methods in Enzymology (Elsevier, 2017; https://linkinghub.elsevier.com/retrieve/pii/S0076687917301143)vol. 591, pp. 55–82.

69. M. Paes Dias, V. Tripathi, I. Van Der Heijden, K. Cong, E.-M. Manolika, J. Bhin, E. Gogola, P. Galanos, S. Annunziato, C. Lieftink, M. Andújar-Sánchez, S. Chakrabarty, G. C. M. Smith, M. Van De Ven, R. L. Beijersbergen, J. Bartkova, S. Rottenberg, S. Cantor, J. Bartek, A. Ray Chaudhuri, J. Jonkers, Loss of nuclear DNA ligase III reverts PARP inhibitor resistance in BRCA1/53BP1 double-deficient cells by exposing ssDNA gaps. Mol. Cell 81, 4692–4708.e9 (2021).

70. A. Meroni, S. E. Wells, C. Fonseca, A. Ray Chaudhuri, K. W. Caldecott, A. Vindigni, DNA combing versus DNA spreading and the separation of sister chromatids. J. Cell Biol. 223, e202305082 (2024).

